# Ratiometric fluorescence nanoscopy and lifetime imaging of novel Nile Red analogs for analysis of membrane packing in living cells

**DOI:** 10.1101/2024.06.03.597272

**Authors:** Line Lauritsen, Maria Szomek, Mick Hornum, Peter Reinholdt, Jacob Kongsted, Poul Nielsen, Jonathan R. Brewer, Daniel Wüstner

## Abstract

Subcellular membranes have complex lipid and protein compositions, which give rise to organelle-specific membrane packing, fluidity, and permeability. Due to its exquisite solvent sensitivity, the lipophilic fluorescence dye Nile Red has been used extensively to study membrane packing and polarity. Further improvement of Nile Red can be achieved by introducing electron-donating or withdrawing functional groups. Here, we compare the potential of derivatives of Nile Red with such functional substitutions for super-resolution fluorescence microscopy of lipid packing in model membranes and living cells. All studied Nile Red derivatives exhibit cholesterol-dependent fluorescence changes in model membranes, as shown by spectrally resolved stimulated emission depletion (STED) microscopy. STED imaging of Nile Red probes in cells reveals lower membrane packing in fibroblasts from healthy subjects compared to those from patients suffering from Niemann Pick type C1 (NPC1) disease, a lysosomal storage disorder with accumulation of cholesterol and sphingolipids in late endosomes and lysosomes. We also find small but consistent changes in the fluorescence lifetime of the Nile Red derivatives in NPC1 cells, suggesting altered hydrogen-bonding capacity in their membranes. All Nile Red derivatives are essentially non-fluorescent in water but increase their brightness in membranes, allowing for their use in MIN-FLUX single molecule tracking experiments. Our study uncovers the potential of Nile Red probes with functional substitutions for nanoscopic membrane imaging.

## Introduction

Cellular membranes are very complex containing a variety of proteins and several thousand different lipid species(1, 2). Phospholipids, sphingolipids, and sterols not only form the structural matrix of cell membranes but also fulfill important signaling functions(3). Their unique properties control membrane permeability, curvature, and packing, thereby providing specific biochemical environments at all types of sub-cellular organelles(4). Membranes of many organelles like the endoplasmic reticulum, Golgi apparatus, or lysosomes are constantly remodeled, requiring recruitment of peripheral proteins, which sense alterations in lipid packing, surface charge, membrane curvature, or viscosity(4). Monitoring such changes in living cells is challenging, and large effort is put into the development of optical probes, which al-low for inferring such membrane properties by optical microscopy. For example, membrane viscosity has been studied by lifetime imaging of molecular rotors, while alterations of dipole and surface potential can be followed by dyes sensitive to changes in the local electric field(5–7).

The packing of lipid bilayers, which biological membranes consist of, is a direct consequence of the hydrophobic effect and can be defined by the extent to which water and small polar molecules can access the bilayer interior. An increase in lipid packing corresponds to a smaller distance between the lipid constituents of the membrane due to a decrease in the area each lipid occupies. This decrease in membrane area due to tighter packing of the lipids restricts the conformational freedom of each lipid. It is often accompanied by a straightening of the lipid acyl chains. Consequently, the membrane thickness increases, while the total bilayer volume remains largely constant(9). The extent of lipid packing in membranes is determined by the acyl chain length, the degree of saturation of phospholipids, as well as by their head group size and charge, and is related to the phase transition temperature (Tm) of a given lipid species (1, 10, 11). At temperatures below the Tm, the membrane lipids form a gel phase with high structural order and slow diffusion. Above the Tm the lipids form a fluid membrane phase, characterized by an increase in their conformational entropy, wobbling, and more rapid diffusion, resulting in decreased membrane packing (10). When a double-bond is introduced in a lipid tail, this will introduce a kink in the chain, demanding more space for acyl chain rotation, which in turn will result in a more loosely packed membrane(11). Cholesterol disrupts the gel phase and orders the fluid phase in a concentration-dependent manner, which often results in a less-sharp (less cooperative) phase transition of lipids in the presence of cholesterol(12). At high cholesterol mole fraction, the biologically relevant liquid-ordered (Lo) phase can form, being characterized by high acyl chain order (tighter packing) but rapid lateral and rotational diffusion of lipids (13). This ability of cholesterol to order fatty acyl chains in the fluid phase and thereby to condense the membrane (i.e., increase its lateral packing) reduces the bending flexibility and is also responsible for the strongly reduced permeability of fluid lipid membranes in the presence of cholesterol (13, 14). Likely for that reason, the plasma membrane (PM) contains the majority of cholesterol, while internal membranes, such as that of the endoplasmic reticulum have to be more flexible and less densely packed, therefore they contain often less cholesterol and more unsaturated phospholipids (15). The picture is even more complicated, since cholesterol-induced changes of membrane properties strongly depend on the acyl chain saturation (16) and protein composition (13, 14). Furthermore, lipid trafficking by vesicular and non-vesicular mechanisms as well as lipid synthesis constantly change the membrane composition in cells(17). Together, this makes understanding lipid organization in cellular membranes particularly challenging.

Membrane packing is closely related to the ability of water to penetrate the bilayer, and a range of dyes have been developed that can monitor the hydration of membranes with varying compositions including hydroxyflavones, Laurdan, and Nile Red(18–20). The latter two probes have been used extensively in cell biology due to their pronounced solvatochromism (19–23). This phenomenon is found in dyes, which experience large changes in the molecular dipole moment during excitation. When such dyes are dissolved in a polar solvent, excitation will generate a reorientation of the polar solvent molecules and this effect will give rise to a red-shifted emission. In contrast, in an apolar solvent, no relaxation of solvent molecules takes place upon excitation of Nile Red, and the emission of the dye will be shifted to the green. Due to these properties and its lipophilic nature, Nile Red is a well-known marker for lipid droplets (LDs)(23).

Particularly in the core of LDs, Nile Red shows a pronounced green emission, which has been used to detect differences in droplet composition and sterol content (20, 23–26). To confine Nile Red to the outer leaflet of the PM, it has been attached to an amphiphilic anchor without perturbing the solvatochromic properties of the dye (27). Using this dye, named NR12S, the authors observed a pronounced color change during endocytosis in a subsequent study, suggesting that lipid packing is lower in endosomes and lysosomes compared to the PM (28). By covalently linking Nile Red to organelle targeting groups, membrane packing could be recently compared between various subcellular membranes (29). By modulating cellular cholesterol content or exerting oxidative and mechanical stresses, organelle-specific responses in membrane packing were observed (29). The pronounced solvatochromism of Nile Red has even been used at the tissue level, in which lipid packing in myelin sheaths from healthy rodents was compared to that of animals with cuprizone-induced demyelination and in postmortem human multiple sclerosis brain(30).

Another intriguing feature of Nile Red is its almost complete absence of fluorescence in aqueous solution, a property that can be used for photoswitching-based super-resolution microscopy of membranes (31). In this method named points accumulation for imaging in nanoscale topography (PAINT), rapid binding of Nile Red to membranes followed by reversible release causes fluorescence blinking which can be used for sparse localization of individual molecules with high precision. The method has been recently combined with spectral imaging, further increasing the sensitivity and resolution in measuring lipid packing in the PM using Nile Red (32). The same principle of rapid reversible binding of Nile Red derivatives to cellular membranes can also be employed in stimulated emission depletion (STED) nanoscopy. Here, photobleached dye molecules can be rapidly replenished with fresh fluorophores, which do not need to be washed out, since Nile Red derivatives are non-fluorescent in the medium and cytosol(33, 34). This enables long-term STED imaging with improved contrast and resolution(34).

To fully explore the potential of Nile Red-based probes, we have recently introduced novel derivatives with strategically placed push-pull groups to increase the solvatochromic responsiveness and other photophysical properties(8). We found that introducing cyano or hydroxy groups at the east side of the molecule can increase the one– and two-photon absorption cross-section. On the other hand, adding a fluorine atom at carbon 10, i.e., at the electron-donating west side of Nile Red, resulted in an analog with increased sensitivity of the fluorescence lifetime towards solvatochromism(8). Introducing electron-donating or ‘push’ groups, like a hydroxy group, or electron-withdrawing or ‘pull’ groups, such as a cyanide group into dye molecules is a widely used strategy to improve their fluorescence properties and to enhance desired attributes, such as their sensitivity to solvent polarity (35–38). However, improved properties in solvents do not necessarily translate directly into better photophysical features in model and cell membranes. Therefore, we have here used selected analogs of Nile Red from our previous study and assessed their potential to image membrane packing in liposomes and in cells. We have used human fibroblasts from healthy donors and from patients suffering from Niemann Pick type C1 (NPC1) disease, a lysosomal storage disorder which is characterized by an accumulation of cholesterol and sphingolipids in late endosomes and lysosomes (LE/LYSs)(39). Using ratiometric STED nanoscopy and fluorescence lifetime imaging (FLIM), we determine the potential of these novel Nile Red derivatives to report about membrane packing in liposomes and living cells. We show that all studied probes can reveal cholesterol-dependent changes in membrane packing, and find that cells lacking functional NPC1 have increasing membrane packing in their LE/LYSs and LDs compared to control fibroblasts. In addition, by employing a pixel-wise principal component analysis (PCA) of FLIM measurements in cells we find that particularly Nile Red senses differences in membrane hydrogen bonding in NPC1 disease compared to control cells. Finally, we employ MINFLUX fluorescence nanoscopy to show, that selected Nile Red probes can be employed for single molecule tracking. Together, our study not only establishes the novel Nile Red derivatives as exquisite sensors of membrane properties but also provides novel insight into lipid packing differences between healthy cells and cells from patients with lysosomal storage diseases.

## Results

### Nile Red derivatives have cholesterol-dependent solvatochromic shifts in membranes and can be imaged by STED nanoscopy

From our library of recently developed Nile Red analogs(8), we decided to use analogs with distinct chemical substitutions in comparison to the parent Nile Red molecule (see Fig. 1a data from Hornum et al.(8)). The rationale behind this choice was foremost to determine the impact of different push-pull modifications on the optical performance of Nile Red analogs in model and cell membranes. In particular, we chose a Nile Red derivative with a ‘push’-type substituent, a hydroxy group at atom 2 (named NR9 here, and analog 7a in Hornum et al.(8)), which donates electrons to the conjugated system, thereby stabilizing the excited state and affecting solvent interactions. In addition, we chose two Nile Red derivatives with ‘pull’-type substitutions, i.e. a fluorine atom at carbon 10 (named NR10 here, and 7n in Hornum et al.(8)). This substitution was speculated to be useful for high-contrast imaging of polar and non-polar interfaces using FLIM (8). Finally, we used a cyano substituted analog at carbon 3 (named NR13 here, and analog 7f in Hornum et al.(8)), which was shown to have an increased molar extinction coefficient and thereby an increased molecular brightness in various solvents compared to Nile Red (see Table S1 and Hornum et al.(8)). These probes have so far only been tested in solvents of different polarities to investigate how this influences their spectral shifts. Fig. 1a shows the recorded spectra measured in Hornum et al.(8), and the extinction coefficient, quantum yield, excitation, and emission maxima in different solvents and can be found in Table S1 as reported by Hornum et al.(8)). Here, we are testing them in a more biological setting of liposomes and live cells (Fig. 1b). While we find that NR9 and NR13 give less fluorescence intensity, NR10 is considerably brighter than Nile Red in liposomes (Fig. 1c-d). Here, it should be noted that excitation is carried out with 561 nm and the emission is collected from 580-754 nm, so the brightness of the liposomes is referred to as the collected photons in this range. From Fig. 1a, S1, S2, and Tab. S1, it can be seen that the behavior of the spectra, quantum yield, and extinction coefficient are very dependent on the local environment. Thus, the conclusion that NR10 is of higher brightness compared to the other Nile Red derivatives is drawn based on the chosen microscope settings and does not resemble an absolute brightness. A similar trend, for Nile Red and the analogs, is observed in human fibroblasts (Fig. 1e), however, here the brightness of NR10 is comparable to the one of Nile Red in the red channel (excitation at 561 nm and emission from 570-720 nm) (Fig. 1e and g). Interestingly, the relative emission in green (excitation at 488 nm and emission from 498 to 551 nm) is higher for NR9, NR10, and NR13 compared to Nile Red (Fig. 1f). The red channel is chosen since this is where most of the fluorescence emission is found (Fig. S1d and e). The green channel is chosen as well, since the amount of the emission found in the green range is increasing with decreasing polarity (Fig. 1a and S1c), so if there are compartments in cells that are even more apolar than toluene, then the emission would be stronger in the green channel. The green emission of Nile Red is often used to characterize neutral lipid deposits, such as LDs in cells(20, 23). The environmental-induced changes in their spectral parameters (excitation, emission, and brightness see Tab. S1 and Fig. S1 and S2) are the main drivers of this effect for all Nile Red derivatives (Fig. 1a). NR9 and NR10 have more green-shifted excitation spectra compared to NR13 and Nile Red (Fig. S1a), which might be the cause for the larger emission in the green channel when incorporated in cells compared to Nile Red. Thus, the green/red emission ratio as defined in our microscope settings, is higher for the substituted Nile Red analogs (Fig. 1g). We note that the exact definition of this emission ratio depends on the chosen detection channels on the microscope, and spectral imaging would be required to determine the full environmental sensitivity of the dyes. Nevertheless, the chosen emission ratio reports sensitively about the local polarity of the membrane environment, in which the Nile Red analogs reside.

**Fig. 1.**
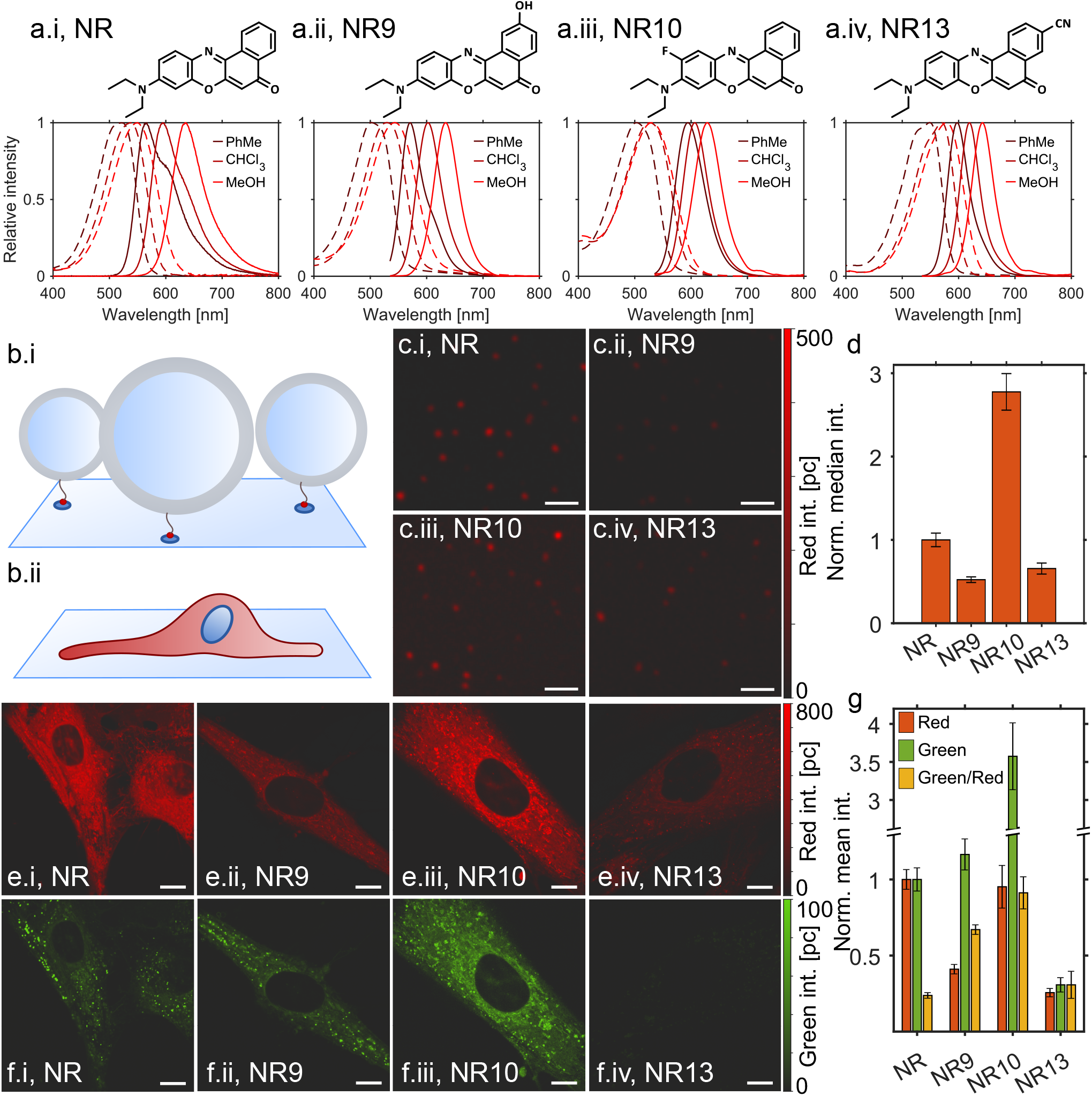
Confocal fluorescence images of Nile Red, NR9, NR10, and NR13 incorporated in vesicles and living ctrl fibroblasts. (a) Shows structures and spectra of Nile Red (a.i), NR9 (a.ii), NR10 (a.iii), and NR13 (a.iv) in toluene (PhMe), chloroform (CHCl_3_), and methanol (MeOH), modified from Hornum et al.(8). (b) Cartoon illustrating experimental setup; vesicles immobilized at a glass surface using a biotin/avidin bond (b.i), and a live cell on a glass slide (b.ii). (c) Confocal fluorescence images of 80 % POPC and 20 % cholesterol vesicles attached to a glass surface intensity adjusted to the same values. Scale bar 2 *µ*m. All images are shown on the same color scale as seen to the right. (d) Bar chart comparing the median fluorescence intensity extracted from the vesicles normalized to Nile Red (number of vesicles for Nile Red: 332, NR9: 233, NR10: 133, and NR13: 54). (e-f) Confocal fluorescence images of live ctrl fibroblasts showing the red (e) and the green (f) emission for cells labeled with Nile Red (e.i and f.i) and the three analogs NR9 (e.ii and f.ii), NR10 (e.iii and f.iii), and NR13 (e.iv and f.iv). All scale bars 10 µm. All images are shown on the same color scale as shown to the right. (g) The bar chart shows a comparison of the obtained mean fluorescence intensity of cells labeled with Nile Red and the three analogs. The intensity was normalized to the mean fluorescence of Nile Red in cells, with the error bars reflecting the SEM between 7 areas of each sample.

Due to its extreme brightness contrast with high fluorescence in membranes and essential absence of emission in water, Nile Red has been used in various forms of super-resolution microscopy including STED(31–33). In STED, the rapid replenishment of imaged and eventually bleached dye in the membrane by dye in the culture medium and cytosol allowed for extended super-resolution imaging(33). We can confirm these observations for all Nile Red analogs, revealing improved resolution by STED in liposomes (Fig. 2a-f). The minimum of the full width at half maximum (FWHM) of the fluorescent liposomes prepared with a nominal diameter of 100 nm was much smaller for STED (FWHM = 71-104 nm) compared to confocal microscopy (FWHM = 186-258 nm). In cells, the lumen of small vesicles in the cytoplasm and the cristae structure of mitochondria could be resolved by STED but not by confocal microscopy for Nile Red and all of its derivatives (Fig. 2g-r). Based on these findings, we employed STED to resolve local solvatochromic shifts of Nile Red and its analogs in human fibroblasts. For that, we collected photons using a 561 nm excitation laser in two detectors; from 640 to 754 nm in detector 1 (det1) and from 580 to 640 nm in detector 2 (det2). The fluorescence ratio det1/det2 is proportional to the polarity-dependent red shift of probe emission. Importantly, all Nile Red analogs change their emission in dependence of several solvent properties, including the dielectric constant, dipole moment, polarity index, and viscosity (Fig. S2). Accordingly, in the complex and anisotropic membrane environment the observed emission ratio of the Nile Red derivatives will report about an interplay of several factors contributing to the overall local polarity. In the membrane, the local polarity changes are caused by differential access of water into the bilayer. To capture the resulting changes in probe emission, we chose detection windows optimized to record solvent-induced emission shifts. In particular, the detector ratio det1/det2 will decrease when the Nile Red analogs reside in a more apolar environment, causing a green shift in their emission (Fig. S1f). Thus, tightly packed membranes with low water accessibility and thereby low contributions of solvent-relaxation to the emission state will have low ratios, while less packed membranes with more water accessibility to the bilayer will have higher det1/det2 values. We found by spectrally resolved STED that this emission ratio is sensitive to the cholesterol content in liposomes for all four Nile Red probes (Fig. S3). Accordingly, all four Nile Red derivatives can faithfully report cholesterol-induced membrane condensation, and the normalized intensity shows the same tendency for STED and confocal measurements of the liposomes (Fig. S3). These results extend earlier studies in model membranes which have shown that the fluorescence emission maximum, intensity, polarization, and lifetime of Nile Red are all sensitive to changes in the cholesterol content of the bilayer(40, 41). Our results demonstrate that this important property of Nile Red is preserved in the functionally substituted Nile Red derivatives.

**Fig. 2.**
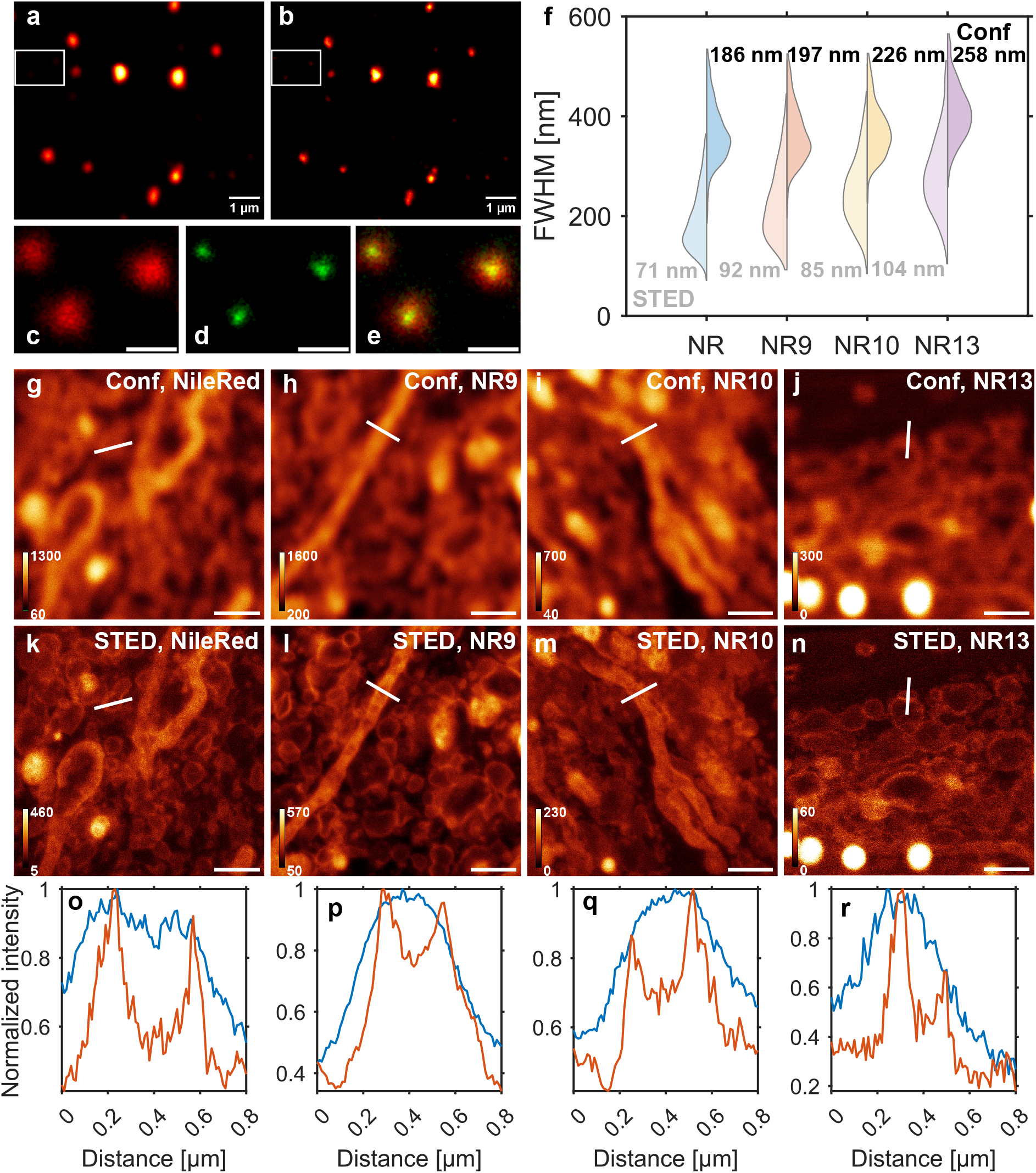
Nile Red, NR9, NR10, and NR13 are all compatible with STED imaging using a 775 nm depletion laser. Comparison of confocal (a) and STED (b) images of vesicles containing Nile Red recorded under PAINT conditions. The smaller zoom at the bottom shows three vesicles significantly smaller than the resolution limit of confocal microscopy (c). The improvement of resolution by STED (d) can be appreciated and is also visible in the overlay (e). Scale bar 500 nm. (f) Double-sided violin plot comparing the FWHM of single vesicles fitted with 2D Gaussians. The FWHM obtained from STED is shown as the distribution to the left, whereas the FWHM obtained from fits to the confocal data is shown to the right. The smallest obtained FWHM is shown in the plot (grey for STED, and black for confocal). Number of vesicles for Nile Red: 1442, NR9: 657, NR10: 470, and NR13: 161. Nile Red derivatives are STED-able to a similar extent as Nile Red in fixed control fibroblasts (g-r). Representative confocal and STED images of fixed cells labeled either with Nile Red (g and k), NR9 (h and l), NR10 (i and m), or NR13 (j and n). Confocal and STED intensity profiles normalized to the maximum value extracted from the white line are compared for Nile Red (o, FWHM *≈* 79 nm), NR9 (p, FWHM *≈* 148 nm), NR10 (q, FWHM *≈* 94 nm), and NR13 (r, FWHM ≈ 100 nm). Color bars on g-n are all shown in photon counts (pc), and all scale bars are 1 µm.

### Organelle specific changes in fluorescence emission ratio of Nile Red and its derivatives in fibroblasts

Using spectrally resolved STED in human control fibroblasts, we first assessed the ability of our experimental setup to detect organelle-specific spectral changes of Nile Red. For that, intracellular membranes captured in detector 1 and detector 2 are shown in Fig. 3a and Fig. 3b, and the green channel used for LD segmentation is shown in Fig. 3c. Local changes in membrane packing are visualized as the ratio between detector 1 and detector 2 (Fig. 3d). Similar images are captured covering different parts of the cells, and an example of an imaged PM with a filopodium is shown in Fig. 3e, f, g, and h. Dividing the imaged cellular structures into five different membrane compartments reveals the lowest emission ratio in the PM, particularly in filopodia indicating the tightest membrane packing (Fig. 3d, h, and i). While this could potentially be a consequence of increased cholesterol content in filopodia, studies with various fluorescent cholesterol analogs do not indicate cholesterol enrichment in particular surface compartments including filopodia (see references in(42)). Instead, altered membrane tension due to strong adhesion between the PM and the underlying cytoskeleton could contribute to the lower emission ratio of Nile Red in filopodia, similar as previously shown for the polarity-sensitive dye Laurdan(43, 44). LDs show a pronounced green emission of Nile Red, which allows for their straightforward identification (Fig. 3c, g, i). We confirmed that by co-staining cells with a blue-fluorescent droplet marker, diphenylhexatriene (DPH), which showed a strong overlap with the green emission of Nile Red (Fig. S5). LDs also exhibit a red emission, which is primarily confined to the droplet-limiting monolayer, as we and others demonstrated for adipocytes and macrophages, where LDs can become very large (24–26). Free cholesterol is known to become enriched in the monolayer surrounding LDs, while cholesteryl esters can be stored in the droplet core(26, 45, 46). Consequently, the det1/det2 red emission ratio of Nile Red in LDs will report the packing of the droplet-limiting phospholipid monolayer, while its green emission is a measure of the size of the hydrophobic droplet core (Fig. 3e-g and i).

**Fig. 3.**
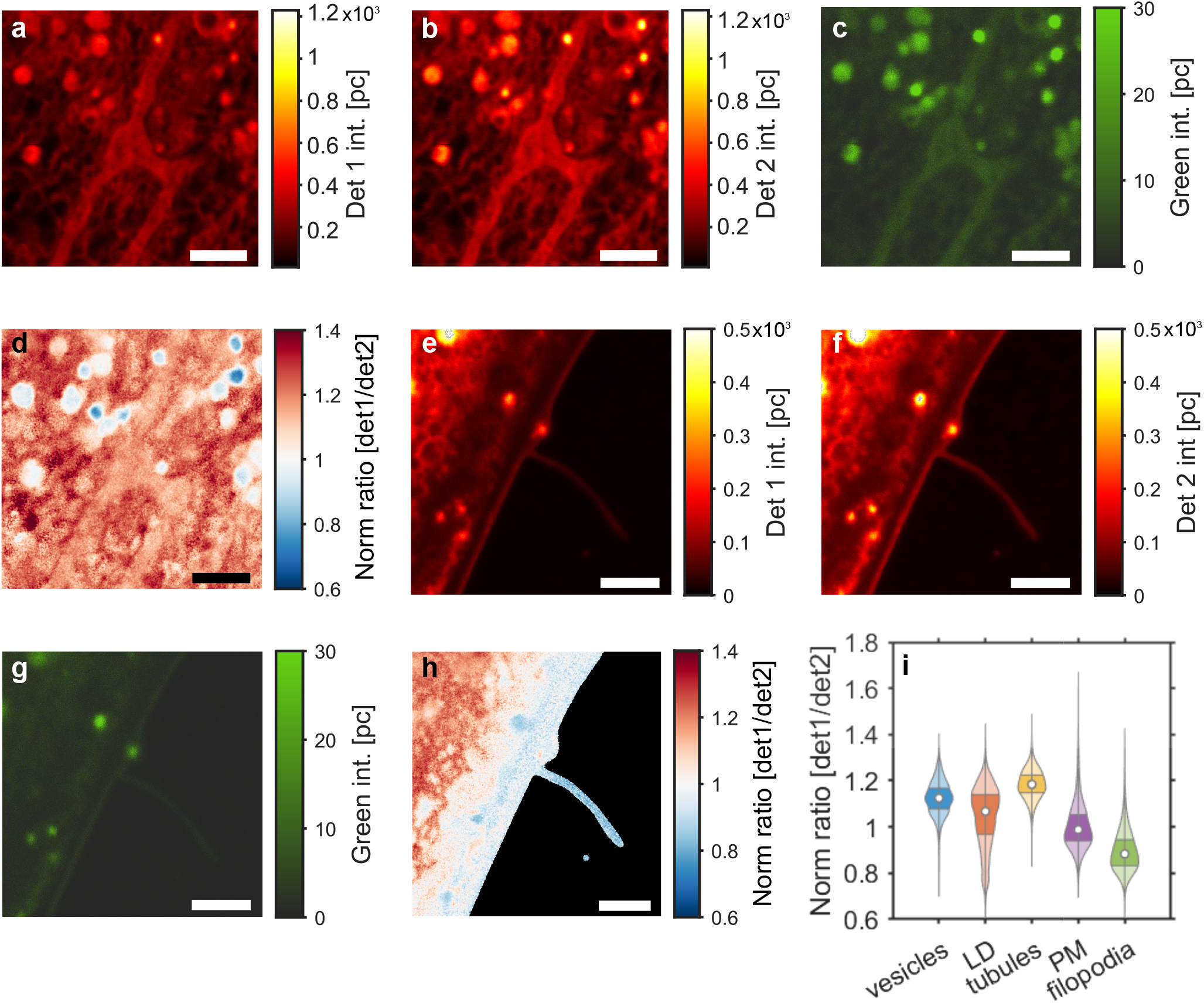
Spectral STED microscopy of membrane packing in living control fibroblasts. STED image of Nile Red emission in detector 1 from 640-754 nm (a) and detector 2 from 580-640 nm (b) of intracellular membranes. (c) the confocal image in the green channel of the area imaged in a and b. (d) shows the normalized ratio (det1/det2) for the area shown in a, b, and c, revealing a distinct lipid packing for different membrane compartments. (e) and (f) show a STED image close to a cell edge from detector 1 and detector 2, respectively. (g) shows the same area as in e and f but for the green channel. (h) shows the normalized ratio of f and e (det1/det2), in which lower values indicate tighter membrane packing. (i) shows a violin plot where normalized pixel ratios between all cells are divided into 5 different subcellular compartments: Vesicles, LDs, tubules, PM, and filopodia. All scale bars are 2 µm.

Mammalian cells shed extracellular vesicles from their PM, particularly during cell migration, signaling, and cholesterol efflux(47). Using a combination of soft X-ray and fluorescence microscopy, we have recently shown that such ectosomes derived from human fibroblasts contain cholesterol markers and sometimes harbor internal vesicles of endolysosomal origin(48). Using STED imaging of Nile Red we also observe ectosomes shedding from the cell surface (Fig. 4a-c), and we confirm that some of them contain internal vesicles (Fig. 4b and e). By ratiometric STED imaging, we found that some of the internal vesicle structures have a lower emission ratio than the surrounding PM, while individual spots with increased emission ratio are also found (Fig. 4d-f). Together, these results underline the potential of spectrally resolved STED microscopy of Nile Red probes for analysis of PM protrusions and membrane vesiculation.

**Fig. 4.**
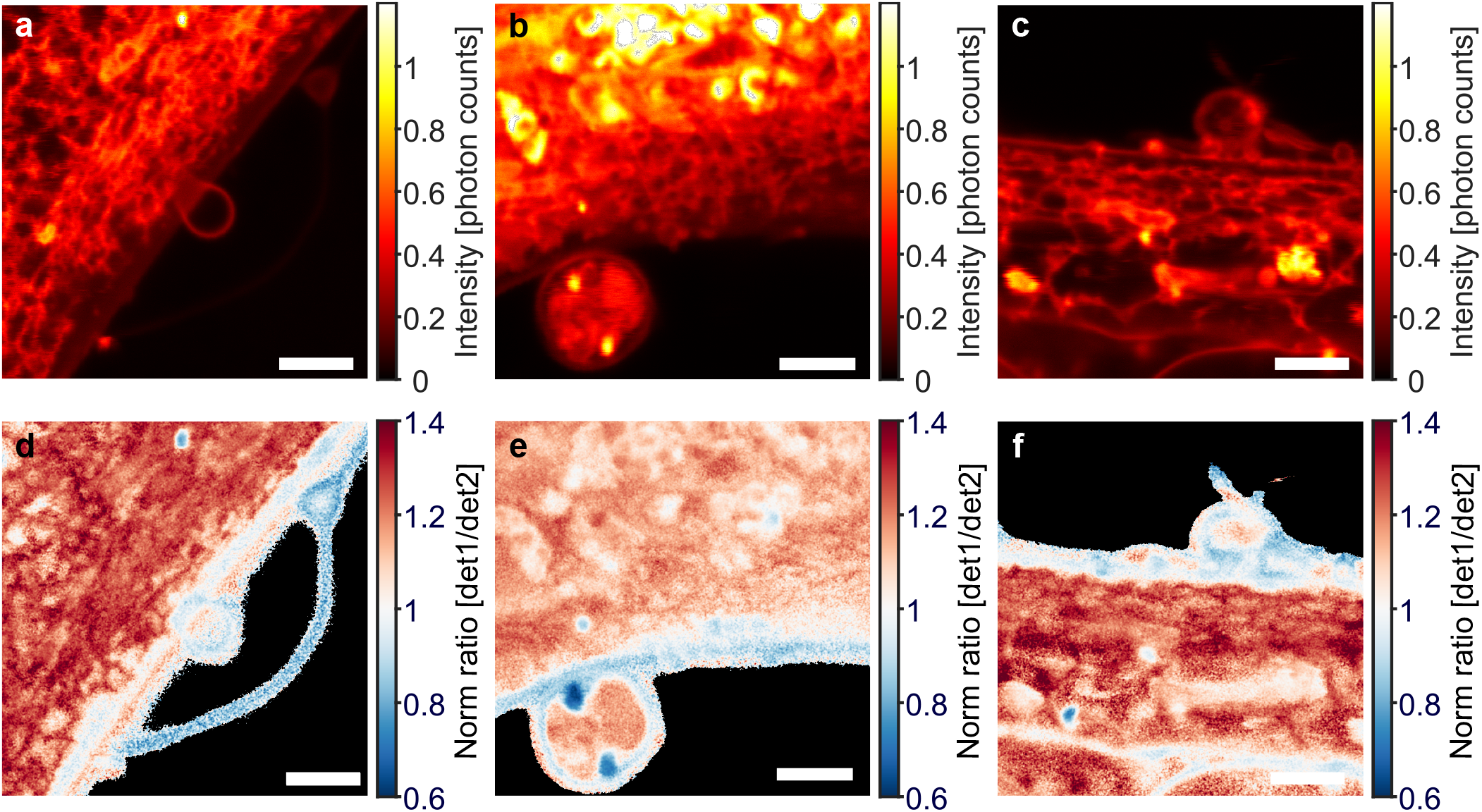
Three different examples of ratiometric STED imaging of ectosomes in control fibroblasts stained with Nile Red. (a-c) show the summed STED images from det1 and det2. (d-f) show the normalized ratiometric STED images of the three positions. Scale bars 2 µm.

Given that Nile Red was also found to stain lamellar lipid phases, as they are found in phospholipidosis and other lysosomal storage disorders(49–51), we decided to compare the Nile Red analogs in fibroblasts from a Niemann Pick C1 disease patient (NPC1 cells) relative to control cells (ctrl cells). In NPC1 cells, the amount of cholesterol and sphingolipids is increased specifically in LE/LYSs due to defective lipid transport out of these organelles(39). Evidence for altered fluidity of the PM in NPC1-deficient fibroblasts was reported, while others only observed alterations in cholesterol-induced membrane packing in internal membranes(52, 53). To determine, whether our novel Nile Red derivatives show different fluorescence properties in subcellular membranes of ctrl and NPC1 cells, we measured the det1/det2 emission ratio in various organelles, which were identified in both cell types (Fig. S6 and Materials and methods). We found that the emission ratio of Nile Red was lower in vesicles and LDs of NPC1-deficient cells compared to ctrl fibroblasts (Fig. 5a). Both NR9 and NR13 had a reduced fluorescence ratio in LDs, while emission of NR10 only showed an obvious change for vesicles (Fig. 5b-d). None of the Nile Red derivatives revealed differences in fluorescence in the PM of ctrl versus NPC1 cells. This result indicates that cholesterol-induced alterations in membrane packing are confined to intracellular membranes in NPC1-deficient fibroblasts, which is in agreement with several studies (53, 54). Other authors reported an increase in lipid order in the PM purified from NPC1 disease compared to control cells, as assessed by fluorescence anisotropy of diphenylhexatriene (DPH) (52). These changes in membrane biophysical properties were paralleled by an increase in saturated lipid species in the membranes of NPC1 cells but a slight decrease in cholesterol content of the PM (52). Adding to this contradiction is the fact, that DPH was added to the cell homogenate in this study, and DPH is known to rapidly redistribute to LDs from the PM (55), where it overlaps with the green emission of Nile Red probes, as we show in Fig. S5. In contrast to the findings by Koike et al. (1998) are results from Somerharju and co-workers, who used high-resolution mass spectrometry to show an increase in unsaturated lipid species in the PM of NPC1-deficient cells, which should lower the lipid packing (56). Thus, our findings are in accordance with the majority of studies concluding that increased lipid packing is confined to intracellular membranes in NPC1-deficient cells and that this is related to the higher cholesterol content of these membranes. Our experiments also show, that different substitutions on the Nile Red core structure result in probes with slightly differing sensitivity to the properties of cellular membranes, an observation, which could be further explored in the future.

**Fig. 5.**
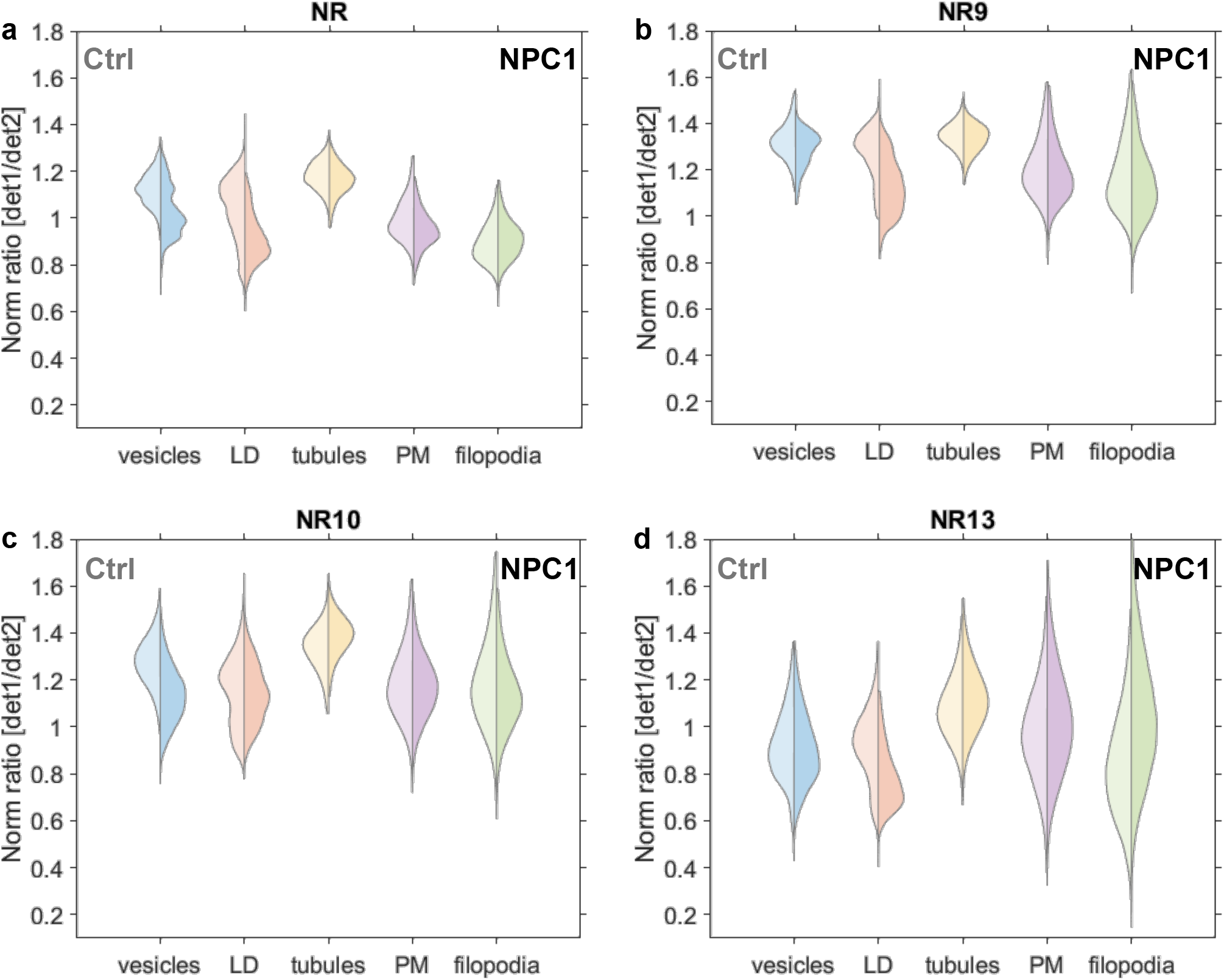
NPC1 cells display a higher membrane packing in vesicles and LDs. Double-sided violin plots display differences in membrane packing between control cells (left) and NPC1 cells (right) for individual sub-cellular compartments; vesicles (blue), LD (red), tubules (yellow), PM (purple), and filopodia (green) for Nile Red (a, N_*ctrl*_: 6, N_*NP C*1_: 6), NR9 (b, N_*ctrl*_: 7, N_*NP C*1_: 7), NR10 (c, N_*ctrl*_: 5, N_*NP C*1_: 7), and NR13 (d, N_*ctrl*_: 5, N_*NP C*1_: 6). Outlier pixels are removed as values scaling more than three median absolute deviations.

### Fluorescence lifetime imaging of Nile Red derivatives in human fibroblasts

To further explore such structure-induced changes in probe sensitivity, we employed FLIM of Nile Red and its derivatives in control and NPC1-deficient fibroblasts. Previous studies have shown that changes in the fluorescence lifetime of Nile Red report about hydrogen bonding capacity of the local probe environment rather than the dielectric solute-solvent interaction(57). Based on these findings we studied the fluorescence lifetime decays of all Nile Red derivatives in cells but found that pixel-wise fitting with exponential functions did not result in any reliable difference between control and disease cells for any of the probes (Fig. S7 and Fig. S8). The reason for this is that the fitting of decay models is sensitive to the signal-to-noise ratio in the images, as we verified in extensive image simulations, where we found that image noise caused a non-linear deviation of the estimated from the true decay constant (Fig. S8). This effect was even enhanced after log-transforming the data, which was necessary to speed up the fitting via linear regression. Apart from noise considerations and computation time, exponential fitting is known not to be unique, which means, one can equally well fit a decay with varying numbers of exponential functions, making the assignment of a given number of components to experimen-tal data difficult(58). This is a well-known but often overlooked limitation, given the fact, that the inner product of real exponential functions is not zero, i.e., they are not orthogonal, in contrast to harmonic functions, used in Fourier analysis. For these reasons, we decided to implement a data-driven method, noise-corrected principal component analysis (NC-PCA), which was recently developed for the analysis of lifetime imaging data(59). Principal component analysis dissects the covariance matrix of any signal into orthogonal components sorted after their contribution to the data variance (i.e., the first principal component is in the direction of the highest data variance, and so on). After noise correction, as done in NC-PCA, the method relies on a singular value decomposition of the data covariance matrix, which allows for detecting orthogonal linear combinations of fluorescence decay components in each data set. Each data point is projected into this eigenvector basis set providing a score image as amplitude for each component(59). NC-PCA applied to fluorescence lifetime data has been shown to detect distinct micro-environments in cell membranes without assuming a pre-defined number of decay components(59). By comparing the score images, corresponding to intensity-weighted principal components, we find that Score 1, representing the contribution with the highest data variation, does not differ significantly between control and NPC1 deficient cells, for none of the Nile Red derivatives (Fig. 6). However, the second component, score 2, resembling the decay component with the second-largest contribution to the total fluorescence decay, is clearly larger in NPC1 deficient compared to control cells, particularly for Nile Red. For NR10, the map of Score 2 revealed differences between intracellular membranes of control and NPC1-deficient fibroblasts. Differences were also found for score 3 and 4, even though these components are relatively small and close to the noise level, making their interpretation more difficult (Fig. S9). Together, these results show, that the fluorescence lifetime of all probes but in particular of Nile Red and NR10, is a sensitive measure for subtle changes in membrane properties of NPC1-deficient cells compared to control cells. We conclude that lifetime measurements can complement ratiometric measurements of Nile Red probes, even though they likely report about other properties of the membrane, such as hydrogen bonding capacity in the bilayer-water interface(57).

**Fig. 6.**
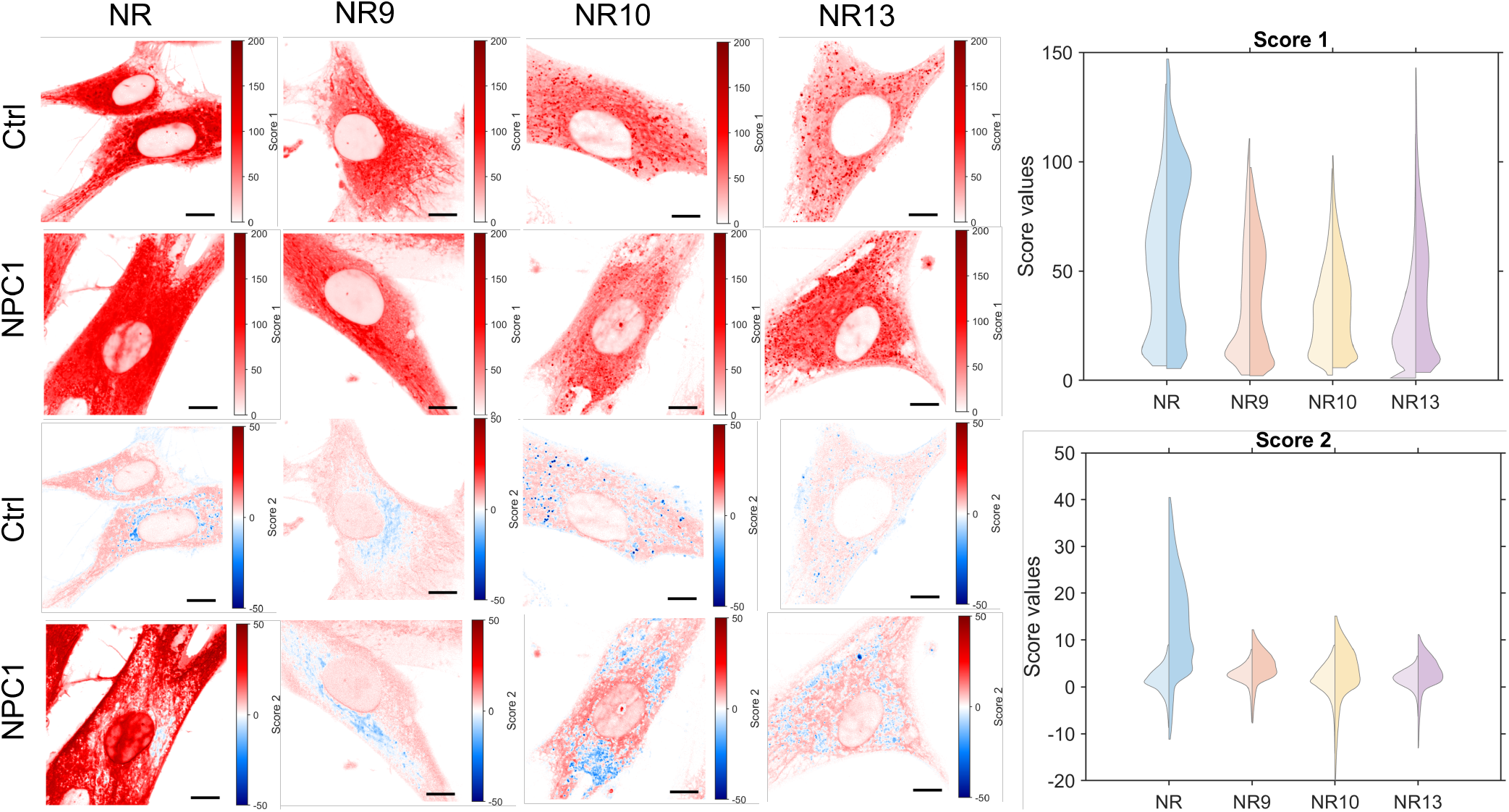
Images for FLIM data generated by NC-PCA analysis for Nile Red, NR9, NR10, and NR13 labeled control and NPC1 cells. Here the estimates for score 1 and 2 are shown as images to the left and collected in double-sided violin plots to the right. The double-sided violin plots compare the distributions of score 1 (top) and score 2 (bottom) for control (left) and NPC1 (right) cells. For each condition, 7 cells are imaged. For score 1 and 2 outlier pixels are removed as values scaling more than three median absolute deviations. Scale bars on the images are 10 µm.

### Single molecule 2D tracking using MINFLUX microscopy

To carry out single-molecule 2D MINFLUX tracking, POPC liposomes were immobilized on a surface and 10 pM of dye was added to the surrounding buffer. The low concentration of dye enables the detection of single molecule binding events to individual liposomes (illustration Fig. 7a) by having one molecule binding a single vesicle at a time. Here, we chose a dye concentration of ≈ 100 times lower than what has previously been reported enabling visualization of single molecule binding of Nile Red to individual vesicles(60) (Fig. S10). Single Nile Red molecule tracks measured in liposomes are shown in Fig. 7b-d, where the liposomes imaged with confocal microscopy are shown in greyscale and the single molecule tracks are overlaid in different colors. To compare Nile Red with the three analogs the length of the individual tracks is collected and shown as a 3D plot (Fig. 7e). Here, it can be seen that NR9, and to a lower extent NR13, has a distribution containing longer trajectories compared to Nile Red and NR10. There can be two reasons for this, 1) NR9 could be more photostable compared to the others, or 2) the release kinetics of NR9 from the liposomes are slower than for Nile Red and the two other analogs. NR10 shows the shortest trajectories of all Nile Red derivatives, which again could be due to rapid release from vesicles and thereby loss of emission or due to faster photobleaching compared to the other Nile Red derivatives. To gain insight into the mobility of Nile Red and the three analogs within liposomes, their speed is calculated over the recorded trajectories. This is done by employing a 35 ms rolling window over the positions and then calculating the speed per position. Nile Red, NR10, and NR13 seem to be moving with an approximate velocity of 1 µm/s, whereas NR9 is moving almost twice as fast (Fig. 7f). The reason for the faster speed of NR9 is unknown at the moment, but one can speculate that NR9 is more aligned with the fatty acyl chains due to the extra hydroxyl group compared to the other Nile Red derivatives, causing its faster lateral diffusion. The extra hydroxyl group in NR9 can act as a hydrogen donor, which could cause a more extensive membrane anchoring and thereby upright membrane orientation of NR9 compared to other Nile Red derivatives. Such an effect has been shown for cholesterol compared to carbonyl cholesterol derivatives, emphasizing the role of hydrogen bonding for membrane packing and dynamics (61, 62). Stronger hydrogen bonding of NR9 could also cause a slower release from the bilayer which in turn might result in an extended on-state and therefore longer single molecule tracks. NR9 also appears to have advantages over Nile Red and the three derivatives when it comes to photon budget (Fig. 7g) and thereby also localization accuracy (Fig. 7h). Since NR9 was found to be less bright in liposomes and cells compared to the other Nile Red probes (Fig. 1c-d), this result obtained with MINFLUX imaging suggests that NR9 might also be more photostable in liposomes than the other analogs. As a consequence of extended single molecule MINFLUX tracking, we found that the mean square displacement (MSD) calculated from tracks of NR9 and NR13 is linear, which is characteristic of Brownian motion (Fig. S11). In contrast, the MSD of Nile Red and NR10 is curved upwards and can be described by an anomalous diffusion model with *α* > 1. Since there is no clear physical model for such a superdiffusive behavior of lipophilic membrane probes, we speculated that this apparent anomalous diffusion is a consequence of poor tracking statistics for Nile Red and NR10. We therefore assessed the impact of time binning and trajectory length on the MSD characteristics (Fig. S12 and S13). We found that varying the time binning of the full-length trajectories when analyzed for the first 1-s interval has almost no effect on the shape of the MSD curve (Fig. S12). In contrast, shortening the trajectories caused superdiffusive behavior also for NR9 and NR13 but to a lesser extent compared to the recordings of NR10 and Nile Red (Fig. S13). We conclude, that a certain trajectory length is needed for reliable MINFLUX analysis. NR9 and NR13 are better suited for single molecule tracking by MINFLUX compared to Nile Red and NR10 because the longer trajectories allow for a more accurate assessment of diffusion properties for NR9 and NR13 in membranes.

**Fig. 7.**
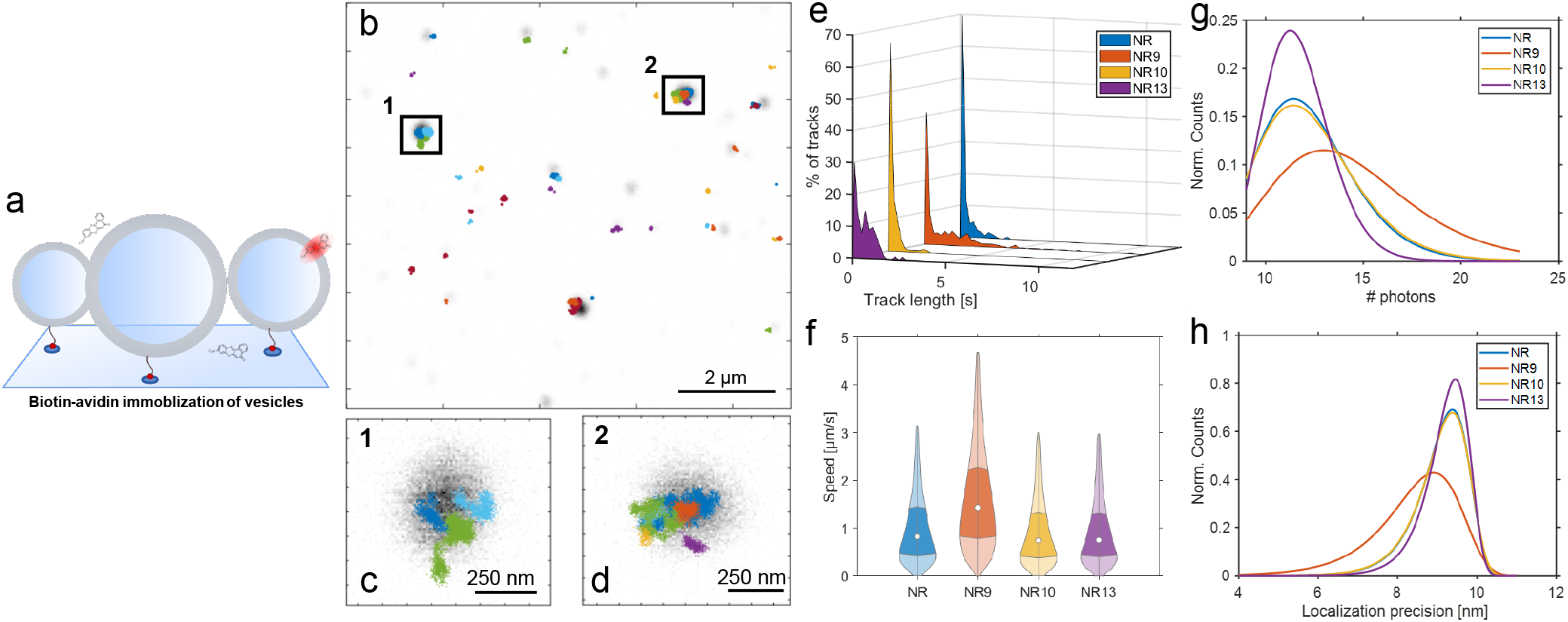
Single molecule MINFLUX 2D tracking of Nile Red and analogs in POPC vesicles. (a) Schematic showing the experimental setup of POPC vesicles immobilized to the glass surface using a biotin-streptavidin system, and Nile Red and the analogs only being fluorescent when incorporated in a vesicle. (b) Overview of POPC vesicles (grayscale) and single molecule 2D MINFLUX tracks of Nile Red shown in the color code. (c-d) Zoom of highlighted areas in (b). (e) Histograms of the track length distributions, normalized to the percentage of tracks. (f) Violin plot showing the speed distributions. Outliers are removed as values scaling more than three median absolute deviations. (g) Log-normal fits to the distribution of the number of photons per localization. (h) Extreme-value distributions fitted to the localization precision of Nile Red and the three analogs. The number of tracks for Nile Red: 158, NR9: 507, NR10: 188, and NR13: 100.

## Discussion

Nile Red is a versatile fluorescent probe, which has been extensively used to stain LDs, cell membranes, myelin, and other lipid assemblies. Due to its pronounced solvatochromism, Nile Red shows significant spectral shifts in LDs versus membranes which has been widely used to identify LDs unequivocally(20, 23–26). More subtle changes in Nile Red’s emission are observed for membranes of different lipid packing, and those can be used to study the unique properties of subcellular membranes with high-resolution microscopy(28, 29, 34). Here, we have studied the potential of novel Nile Red analogs to assess lipid packing in fibroblasts from healthy donors and from NPC1 disease patients using spectrally resolved STED nanoscopy and FLIM. We find that Nile Red and all studied analogs can identify the increased lipid packing of subcellular membranes in NPC1-deficient cells compared to fibroblasts from healthy donors. Cholesterol is well-known to condense membranes consisting of phospholipids in their fluid state, i.e., above the Tm of the phospholipids’ chain (see Introduction section). At high cholesterol mole fraction, particularly in mono-or disaturated phospholipid membranes, this property of cholesterol can induce the Lo phase which has been studied in detail in model membranes. Cellular membranes rich in cholesterol and saturated phospholipids, such as the PM and recycling endosomes have been shown to possess Lo-like properties, and this can explain the pronounced blue-shift in the emis-sion of the Nile Red probes studied here. However, when interpreting emission changes of solvent-sensitive dyes, like Nile Red in terms of cholesterol-induced changes in the physical properties of cell membranes, one has to be careful. This is because the acyl-chain ordering by cholesterol is paralleled by other changes, like altered membrane tension, permeability, fluidity, lateral and rotational diffusion, or bending flexibility of the membranes. While Nile Red probes are sensitive to lipid packing differences, they also respond to changes in rotational motion and membrane tension, making a molecular interpretation of the observed emission changes challenging(41, 63).

We find that the novel Nile Red analogs behave overall similarly to the parent Nile Red molecule concerning their environmental sensitivity. However, there exist small but important differences, which can be used as an advantage for a given application. For example, NR10 bearing a fluorine substituent at the west side of the molecule shows more pronounced variations in FLIM compared to NR9 and NR13 (Fig. 6, score 2), however, Nile Red exhibits the largest dynamic range of score 2. On the other hand, NR9 and to a lower extent NR13, containing an extra hydroxyl and cyanide group on the molecule’s east side, respectively, seem to be better suitable for single-molecule tracking than Nile Red and NR10, even though NR10 is the brightest analog in liposomes (Fig. 1d). The increased brightness of NR10 when incorporated in vesicles and cells is surprising considering the brightness in solvents is the lowest of them all(8). A particularly interesting aspect of NR10 is the absence of emission in polar solvents, but the relative brightness in polar environments, here the brightness almost increases with a factor of 10 from MeOH to CHCl_3_ (Tab. S1)(8). Other solvent properties also seem to affect the emission maximum (Fig. S2), and could also influence the brightness of the molecule, however, these effects have not been tested in this study.

NR9 followed by NR13 give longer single molecule tracks compared to the other Nile Red probes, suggesting that these derivatives are more photostable under conditions used for 2D MINFLUX tracking. Alternatively, NR9 and NR13 could reside longer in the membrane than the other Nile Red derivatives, for example due to slower dissociation from the bilayer. This would allow for longer ‘on-states’ and thereby for prolonged MINFLUX imaging. Longer tracks, in turn, allow for a more precise estimation of diffusion behavior, which is reflected by the observation of normal diffusion in the membrane for NR9 and NR13 but not the other Nile Red derivatives. NR9 also gives the highest number of photon counts, allowing for more accurate determination of molecule positions by MINFLUX imaging. In contrast, Nile Red and NR10 show apparent anomalous subdiffusion behavior, likely because trajectories are too short for obtaining sufficient statistics. These findings show that our strategy of introducing chemical substitutions into the Nile Red structure allows for fine-tuning the fluorescence response of the probes in intra-cellular environments. Due to the possibility of measuring photon counts in two detectors at once, ratiometric tracking studies of Nile Red and analogs, such as NR9, future stud-ies could reveal changes in lipid packing in liposomes or even in cells at the single molecule level. This could grant us a method to resolve areas of different membrane packing in cells and even correlate the emission color with the actual speed of the single molecules. We tried initial single molecule MINFLUX 2D tracking experiments in living cells with Nile Red and NR9 (see Fig. S14). This serves as proof of principle, that MINFLUX tracking can be carried out in living cells, and it suggests that NR9 might be a better dye for these experiments than Nile Red and its other derivatives due to longer track lengths. However, due to the presence of auto-fluorescence in cells, further studies are needed to validate the method, which are beyond the scope of this article. Nile Red and all its studied analogs showed blue-shifted emission in NPC1-deficient cells compared to control cells, particularly in subcellular vesicles and droplets. This indicates higher lipid packing due to the increasing content of cholesterol and other lipids. Also, more subtle differences between the dyes are seen, as only Nile Red and NR10 detect a changed lipid packing in vesicles of NPC1 cells, while all probes except NR10 report altered emission in LDs (Fig. 5). Thus, all Nile Red derivatives can be used as sensitive sensors for changes in lipid packing in lysosomal storage diseases and a contribution of several Nile Red probes can increase the sensitivity of the analysis.

## Methods

### Materials and reagents

Nile Red and all of its derivatives were synthesized as described previously(8). **Chemicals:** 1-palmitoyl-2-oleoyl-glycero-3-phosphocholine (POPC, Avanti, 850457P), Biotinyl Cap PE (DOPE-biotin, Avanti, 870273), Cholesterol (Chol, Sigma, C8667-1G), 1-palmitoyl-2-6-[(7-nitro-2-1,3-benzoxadiazol-4-yl)amino]hexanoyl-sn-glycero-3-phosphocholine (NBD-PC, Avanti, 810130P), Avidin (Sigma, A9275-2MG), 1,6-diphenyl-1,3,5-hexatriene (DPH, Sigma, D208000-1G), Dimethyl Sulfoxide (DMSO, Sigma, D2650-100ML), mini extruder (Avanti, 610020), and 200 nm polycarbonate filters (Whatman™,10417004). **Cell culture:** Human fibroblasts from a healthy male donor (Coriell Institute #GM08680 (referred to as control)) and from a patient with NPC1 disease (Coriell Institute #GM03123) were purchased from Coriell Cell Repositories. Minimum Essential Medium (MEM, Gibco, cat.no.: 42360-032), MEM Non-essential Amino Acid Solution (NEAAS, Sigma, M7145-100mL), Fetal Bovine Serum (FBS, Gibco, 10270-106), Dulbecco’s Modified Eagle’s Medium (DMEM, Sigma, D6429-500MK), phosphate buffered saline (PBS, Gibco, 70013-016), M1 media (150 mM NaCl (Merck, 1.06404.1000), 5 mM KCl (Merck, 104936), 1 mM CaCl_2_ (Merck,2382.1000), 1 mM MgCl_2_ (Merck, 1.05833.1000), 5 mM glucose (Merck, 1.08342.1000), and 20 mM HEPES (Sigma, H3375-100G), pH addjusted to 7.4), penicillin-streptomycin (P/S, Sigma, P4333-100ML), Trypsin-EDTA (Sigma, T4174-100ML), GlutaMAX (Gibco, 35050-038), 35 mm microscope dishes (P35G-1.5-50-C, MatTek), and µ-Slide 8 Well high Glass bottom #1.5H glass coverslip (Ibidi, 80807).

### Liposome preparation

Four different liposome compositions were prepared: 1) 99.5 mol% POPC, 0.5 mol% DOPE-biotin, 2) 79.5 mol% POPC, 0.5 mol% DOPE-biotin, 20 mol% chol, 3) 59.5 mol% POPC, 0.5 mol% DOPE-biotin, 40 mol% chol, and 4) 99 mol% POPC, 0.5 mol% DOPE-biotin, 0.5 mol% NBD-PC. Compositions 1-3 were used to quantify the influence of chol on the emission of Nile Red and the three analogs in STED. Composition 4 was used for MINFLUX single molecule tracking experiments of Nile Red and the three analogs. The lipids for the four different vesicle compositions were thoroughly mixed in glass vials from CHCl_3_ stocks and for the DOPE-biotin from an EtOH stock in the molar ratios stated above. The solvent was evaporated under N_2_-flow, and the lipid film was placed under vacuum for >2 h. The lipid films were rehydrated for 30 min in 1x PBS in a concentration of 1.25 mM and the solution was gently swirled. The newly formed vesicles were exposed to 8 freeze thaw cycles freezing in liquid N_2_ and thawing in a water bath to minimize multilamellarity. The liposomes were then extruded through a 200 nm polycarbonate filter in an Avanti mini extruder. Ibidi 8-well µ-Slides were used for liposome imaging. The chambers were plasma cleaned for 2-5 min at medium strength in an expanded plasma cleaner from Harrick Plasma. Immediately when exiting the chamber of the plasma cleaner 50 µL of 0.05 mg/mL avidin in H_0_O was added to each of the 8 wells, and the chamber was incubated for 40 min at room temperature. Then it was washed three times with PBS and either used immediately or kept at 4°C overnight.

### Cell culture

Both control and NPC1 cells were grown at 37°C with 5 % CO2 under 100 % humidity. The NPC1 fibroblasts were grown in MEM supplemented with 15 % FBS, 1 % P/S, and 1 % NEAAS. The control fibroblasts were grown in DMEM supplemented with 10 % FBS, 1 % GlutaMAX, and 1 % P/S. Before plating both cell lines were flushed once with 1x PBS and trypsinized for 5 min at 37°C with 5 % CO2 under 100 % humidity before they were plated on plain glass on 35 mm microscope dishes. Both cell types were left to settle for 24 hours before imaging. All experiments on cells were carried out following ethical guidelines and safety regulations defined by the provider Coriell Cell Repositories (www.coriell.org) and the University of Southern Denmark.

### Cellular labelling solutions

Before labeling the cells, Nile Red, NR9, NR10, and NR13 were prepared as 1 mM stocks in DMSO. Before labeling the cells were washed once with M1-media. A labeling solution of M1-media and 1.6 µM of Nile Red or one of the analogs was added to the cells and the imaging was started almost immediately after at room temperature. For the colocalization experiment between DPH and the green signal from NileRed. The control fibroblasts were initially stained with DPH, here a 4 µM DPH solution in M1-media was prepared from a 2 mM DMSO stock, and the cells were incubated for 20 min at 37°C with 5 % CO2 and 100 % humidity. The DPH labeling solution was removed and imaging media containing 1.6 µM Nile Red was directly added to the cells without any further washing steps.

### STED microscopy

#### Live cell imaging

STED live cell and vesicle imaging was performed on an Abberior Facility Line STED microscope (Abberior Instruments GmbH). All excitation and STED lasers are pulsed and circularly polarized. Confocal imaging of Nile Red and the analogs was carried out in two-channel a green channel excited at 488 nm and emission collected between 498-551 nm, and a red channel excited at 561 nm and emission collected between 570-720 nm, with a pixel size of 80 nm. Confocal imaging of DPH was carried out using a 405 nm excitation laser, and emission was collected between 415-478 nm.

STED imaging of Nile Red and the analogs were measured in 3 different channels all collected simultaneously. The green channel excited at 488 nm and emission was collected between 498-551 nm. The other two channels were collected in the red and the far-red part of the emission spectrum, here a 561 nm excitation laser was used, and the emission signal was split into two detectors; detector 1 between 640-754 nm, and detector 2 between 580-640 nm. It was possible to use a 775 nm 2D STED laser to increase resolution in the two red channels. Here, 750 ps gating was applied with a measurement width of 8 ns.

The spectra of Nile Red and the three analogs will vary in dependence on the local environment of the dyes (Fig. 1a, S1, S3, and S4). The green detection channel is placed to collect the largest amount of signal between the 488 nm and the 561 nm excitation lasers leaving 10 nm on each side of the detector. 561 nm is chosen as an excitation source for the red STEDable part of the spectrum, which has also been used for Nile Red previously(33, 34). When applying the STED laser we only observe an improved resolution for the red channel, collected in detector 1 and detector 2, but not for the green channel, which has been measured simultaneously. Detector 2 leaves a gap of 10 nm to the 561 nm excitation laser, which is preferred to protect the detector from any excitation light. This choice for detection channels can sensitively detect changes in cholesterol-induced membrane packing (Fig. S3 and S4), and we also observe organelle-specific packing in cells (Fig. 3 and 5).

FLIM imaging was carried out by using a 561 nm pulsed laser, and a repetition rate of 40 MHz. The emission was collected between 571-720 nm, pixel size of 60 nm with a pixel dwell time of 100 µs. For Nile Red photons were collected with an offset of 3 ns for 19.9 ns in a time interval of 0.21 ns, for the analogs a shorter offset of 2 ns was chosen. For all imaging, a 100x magnification UplanSApo 1.4 NA immersion oil objective lens was used.

#### Liposome imaging

Liposome compositions 1-3 were labelled with Nile Red and the three analogs and imaged under the same conditions as the cells. In short 1 µL of the liposome master mixture was added to 990 µL 1xPBS containing 1.5 µL of 1 mM Nile Red, NR9, NR10, or NR13, and then 200 µL of this liposome mixture was added to a well in the avidin coated 8-well Ibidi µ-Slide. The solution was left to settle for ≈ 10 min before image acquisition was started under the same conditions as the live cell imaging.

### MINFLUX 2D tracking

Single-molecule tracking of NileRed, NR9, NR10, and NR13 was carried out in vesicle composition 4, POPC vesicles containing 0.5 mol% DOPE-biotin and 0.5 mol% NBD-PC. Here, avidin-coated 8-well Ibidi µ-Slides were used as well. Before imaging the wells were coated with 100 nm gold beads as well, this was done by adding 100 µL gold beads directly from the stock to each chamber followed by an incubation period of 7 min at RT, before the excess beads were removed and the surface was flushed once with 10 mM MgCl_2_, and then once with 1xPBS, before 100 µL of liposomes diluted 1:500 from the stock was added to the chamber. All MINFLUX measurements were carried out on a commercial MINFLUX microscope from Abberior Instruments GmbH. Here, the NBD signal was measured in confocal-mode excited using a continuous 488 nm laser, and emission was collected between 500-550 nm, and the MINFLUX channel was checked for bleedthrough from the NBD (excitation 560 nm, emission 580-630 nm), where there were found no traces. Then either Nile Red or one of the three analogs were added in a concentration of 10 pM to the chamber, to have the fluorophores dilute enough to be able to distinguish single molecule incorporation events in single vesicles. Small areas containing spacially distributed single vesicles in the NBD channel were selected and then fast 2D MINFLUX tracking was applied using the 560 nm MINFLUX laser with emission detected between 580-630 nm. The data was exported in the .mat format allowing quantification in MATLAB.

### Data analysis

#### Characterization of Nile Red and analogs in cells

The brightness analysis in figure 1 was carried out on confocal images of entire cells. Mean intensity was calculated for each image by omitting all values below a background threshold and then calculating the mean of the remaining pixels. For the red channel, the background intensity was determined from the intensity values where there were no cells. The background intensity for the green channel was determined from the unspecific signal from the cells. The green and red intensity values were normalized to the mean value of the Nile Red intensity in control cells.

#### Colocalization of DPH and Nile Red

Background thresholds were applied to all images by setting the background level to 0. Similarly to the characterization the background for the red channel was determined from the intensity values where there were no cells, and for the green and blue channels, it was determined from the unspecific signal from the cells. The amount of colocalization between DPH, the green, and the red contribution of Nile Red were assessed by the Pearson coefficient(64):

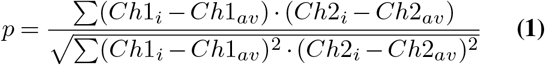

Here, *Ch*1_*i*_ is the pixel intensity of each pixel in channel 1, and *Ch*1_*av*_ is the average intensity of the entire image in channel 1. *Ch*2_*i*_ and *Ch*2_*av*_ are the same values but for the corresponding channel 2.

#### Ratiometric STED measurements

To determine the ratio between detector 1 and detector 2, a Gaussian filter with a sigma of 1.5 was applied to the image of each detector before calculating the ratio between them. To ensure that only specific signals were used in the analysis a background level outside the cells was chosen and only signals above this level were used. The ratiometric measurements were divided into sub-cellular membrane compartments, this division was carried out manually by drawing individual masks on top of the red intensity images for vesicular structure, tubules, the PM, and filopodia. The vesicular structures were further separated into two categories, vesicles and LDs. The LDs were defined based on the intensity of green emission found within the vesicles. The filtered version of Fig. 3f can be seen in Fig. S6b, and the vesicle and tubules mask for this area can be seen in Fig. S6c and d, respectively. All violin plots are generated using the MATLAB code by (65).

#### FLIM analysis

All FLIM traces were linearized taking the natural logarithm of each pixel’s decay curve. Thereby a single exponential could be fitted by a linear function which decreases the computation time significantly, and the life-time could be extracted by one over the slope. The extracted slopes for ctrl and NPC1 cells are plotted in a double-sided violin plot in Fig. S7.

#### Simplified FLIM simulation

To investigate how noise affects the fitting of single exponential lifetime data, we carried out a simple simulation where we simulated a FLIM stack with a single exponential decay function:

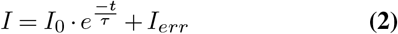

where *I* is the intensity at timepoint *t, I*_0_ is the initial intensity, *τ* is the fluorescence lifetime, and *I*_*err*_ is uniformly distributed around 0 semi-random noise of varying degrees (spanning from 0 to 40 intensity counts) to the system by the use of the randi-function in MATLAB. We then defined 8 individual image patches consisting of 16×16 pixels, where each pixel was generated individually. For each patch, we used an increasing value of *I*_0_ ranging from 20 to 300 in intervals of 40, and *τ* was kept constant at 4 ns. The integrated intensity images are seen in Fig. S8a.i-v. The lifetime for each pixel is then extracted by either fitting a single exponential decay and extracting the lifetime (Fig. S8b.i-v shows the extracted lifetimes) or by linearizing the FLIM stack and fitting with a linear function (Fig. S8c.i-v). For each 16×16 pixel square, we calculated a mean extracted lifetime and standard deviation, for *I*_0_ = 100, 180, and 300, the mean and standard deviation are plotted as a function of the noise in the overall image.

#### Noise-corrected principal component analysis

Noise-corrected principal component analysis (NC-PCA) was carried out as described by LeMarois et al.(59). In short, the image stacks are Poisson noise corrected by dividing each image with the square root of the mean intensity of each time-bin. After the noise correction, a PCA analysis is used to obtain a time-scale separation of individual components. For each obtained principal component a score value for each pixel can be obtained:

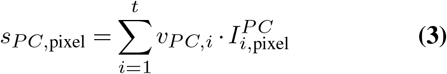

Thereby a score value is obtained for each pixel in an image and denotes the spatial separation of FLIM microdomains in an image.

#### 2D MINFLUX tracking analysis

2D MINFLUX tracking data was loaded in MATLAB R2022b using the loading function, minfluxmat2loc, found in the software from Jonas Ries’ lab at EMBL in Heidelberg(66). Then the data was assigned to individual single molecule tracks and from these track lengths of individual single molecule binding events to the liposomes could be quantified (Fig. 7e). This track length can be aborted for 3 reasons, 1) because the single molecule of Nile Red, NR9, NR10, or NR13 is leaving the vesicle, and the change in brightness upon the change in environment makes it dark, 2) it bleaches, and 3) the track is aborted because a second molecule enters the vesicle. Here, we diluted the concentration of Nile Red and the analogs to an extent so the second molecule entering the vesicle seems unlikely. However, if it is bleaching or unbinding from the vesicle that causes the track to stop we are not sure. To calculate the speed at which the single molecule moved we applied a sliding window of 35 ms(67) to the accumulated distance the molecule had traveled and divided this with the time (Fig. 7f). For each localization in the single molecule tracks there will be collected a certain number of photons. The more photons collected the better localization. Here, the photon distributions followed log-normal distributions, so each distribution from Nile Red, NR9, NR10, and NR13 are fitted to a log-normal for better visualization and they are shown in Fig. 7g. From the number of photons, the localization precision approximated to(66, 67):

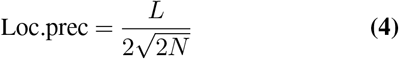

Given that 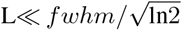, where L is the size of the probing range given in nm, *fwhm* is the full-width-half-max of the beam size, and N is the number of photons. The localization precision histograms are fitted with extreme-value distributions to ease visualization (fig. 7h).

## Acknowledgements

The authors are very grateful for the MATLAB code shared by Dr. A. LeMarois(59). We acknowledge the Independent Research Fund Denmark – Natural Sciences (DFF-FNU) for financial support (Grant ID: DFF-7014-00050B). Image acquisition was performed at the Danish Molecular Biomedical Imaging Center (DaMBIC, University of Southern Denmark), supported by the Novo Nordisk Foundation (NNF) (grant agreement number NNF18SA0032928). The authors acknowledge CellX (The Danish Single Cell Examination Platform) for access to the MINFLUX microscope. DW acknowledges funding from the Lundbeck Foundation (grant no. R366-2021-226) and from the Danish Research Council (grant ID: 2034-00136B).

## Author contributions statement

All authors contributed to the concept and the main idea. D.W., M.S., and L.L. conceived the experiments. L.L. conducted the experiments. L.L. and D.W. analyzed the results and wrote the first draft of the manuscript. M.S., M.H., P.R., J.K., P.N., and J.R.B. reviewed and commented on the manuscript.

## Data availability statement

All data used in this article will be available upon request to the corresponding author.

## Ethics declarations

The authors declare no competing interests.

