## Supplementary Information for "Ratiometric fluorescence nanoscopy and lifetime imaging of novel Nile Red analogs for analysis of membrane packing in living cells"

### Fluorescence sensitivity of Nile Red, NR9, NR10, and NR13 to polar and apolar solvents

Here, the spectral data is obtained from Hornum et al.(8). In table S1 the excitation and emission maximum, quantum yield, and extinction coefficient can be found in solvents of varying polarity, methanol being the most polar and toluene being the most apolar. How the excitation and emission max change dependent on the solvents are plotted in Fig. S1a and b, for all four dyes showing a decrease in both excitation and emission maximum upon decreasing polarity of the solvent.

| Dye | Solvent | Excitation max [nm] | Emission max [nm] | Quantum yield | Extinction coefficient [$mM^{-1}cm^{-1}$] |
|---|---|---|---|---|---|
| NileRed | PhMe | 521 | 566 | 0.83 | 28.9 |
| | $CHCl_3$ | 537 | 594 | 0.64 | 32.6 |
| | MeOH | 550 | 636 | 0.34 | 32.6 |
| NR9 | PhMe | 502 | 571 | 0.56 | 32.3 |
| | $CHCl_3$ | 530 | 603 | 0.44 | 36.1 |
| | MeOH | 544 | 634 | 0.28 | 34.9 |
| NR10 | PhMe | 503 | 595 | 0.52 | 22.9 |
| | $CHCl_3$ | 527 | 607 | 0.45 | 25.2 |
| | MeOH | 530 | 629 | 0.04 | 26.9 |
| NR13 | PhMe | 549 | 598 | 0.49 | 47.4 |
| | $CHCl_3$ | 570 | 619 | 0.49 | 42.6 |
| | MeOH | 578 | 642 | 0.22 | 36.8 |

**Table S1.** Excitation maximum, emission maximum, quantum yield, and extinction coefficient measured in toluene (PhMe), chloroform ($CHCl_3$), and methanol (MeOH) from preceding publication by Hornum et al.(8).

DRAFT

To investigate how these spectral changes influence the results seen in the cells on the microscope, we divided the spectra into the three different channels used at the microscope, the green channel (498-551 nm), and the two red channels, detector 1 (640-720 nm) and detector 2 (580-640 nm). For each channel, we calculated how much of the emission spectrum is found in the respective channel. This is seen for the green channel (Fig. S1c), detector 1 (Fig. S1d), and detector 2 (Fig. S1e) as a function of the solvents. It shows that NR9 has the largest shift of the four dyes, moving from close to 0 % of the emission spectrum in the green in MeOH to close to 25 % of the emission spectrum in the green when changing the solvent to the less polar toluene (Fig. S1c), making the green shift of NR9 the most sensitive of the four dyes. For detector 1, all four dyes show a green shift resulting in less percentage of the spectrum being in detector 1 upon decreasing the polarity of the solvent (Fig. S1d). However, in detector 2 (Fig. S1e), an increase in spectral percentage and then a decrease is seen for Nile Red and NR13, indicating the spectrum moving through the bandpass of the detector, whereas it is almost constant for NR10 and decreasing for NR9. When plotting the ratio between detector 1 and detector 2 (Fig. S1f) dependent on solvent, we observe a decrease in ratio with a reduction of solvent polarity. This is the same overall trend as seen for liposomes with increasing cholesterol content (Fig. S3). Here, the largest reduction in the ratio is seen for Nile Red and NR13, indicating that these two dyes by this measure might be the most sensitive to environmental changes. However, it should be noted that the raw spectral ratios (det1/det2) for $CHCl_3$ and PhMe are significantly smaller than the raw emission ratios in solvents observed in cells and liposomes (Fig. S4), suggesting that these two solvents have a lower polarity than what is found in cellular membranes and liposomes. In support of this analysis, we also found that all Nile Red derivatives are similarly sensitive to cholesterol-induced changes in lipid packing in liposomes (see below and Fig. S3).

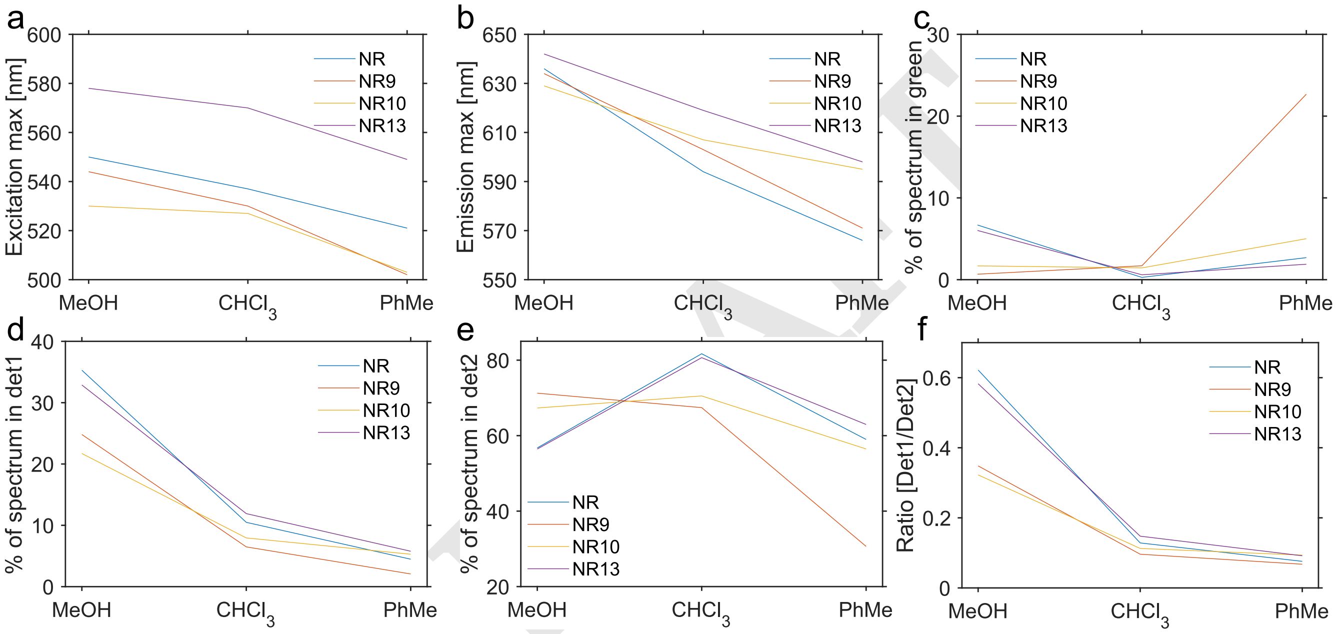


**Fig. S1.** Spectral characterization of Nile Red, NR9, NR10, and NR13 in methanol (MeOH), cholorform ($CHCl_3$), and toluene (PhMe). (a) and (b) show a graphical representation of the excitation and emission max in the three different solvents, respectively. (c), (d) and (e) show the percentage of the spectra in the green, and two red channels (detector 1 and detector 2) used for confocal and STED imaging dependent on the different solvents. (f) shows how the ratio of integrated intensity in detector 1 over detector 2 varies depending on the different solvents. The original data of the spectra obtained from Hornum et al.(8).

***Solvent properties influence on emission maximum.*** Since we haven't clearly defined solvent polarity in this study, we want to show that different solvent properties can influence the emission maximum of Nile Red and the three analogs. Here, we have plotted four properties; the dielectric constant(68) (Fig. S2a), the dipole moment(68) (Fig. S2b), the polarity index(68) (Fig. S2c), and the viscosity(69) (Fig. S2d) against the emission maximum of Nile Red and the three analogs. We find that all these solvent parameters affect the spectral shift of Nile Red probes (Fig. S2).

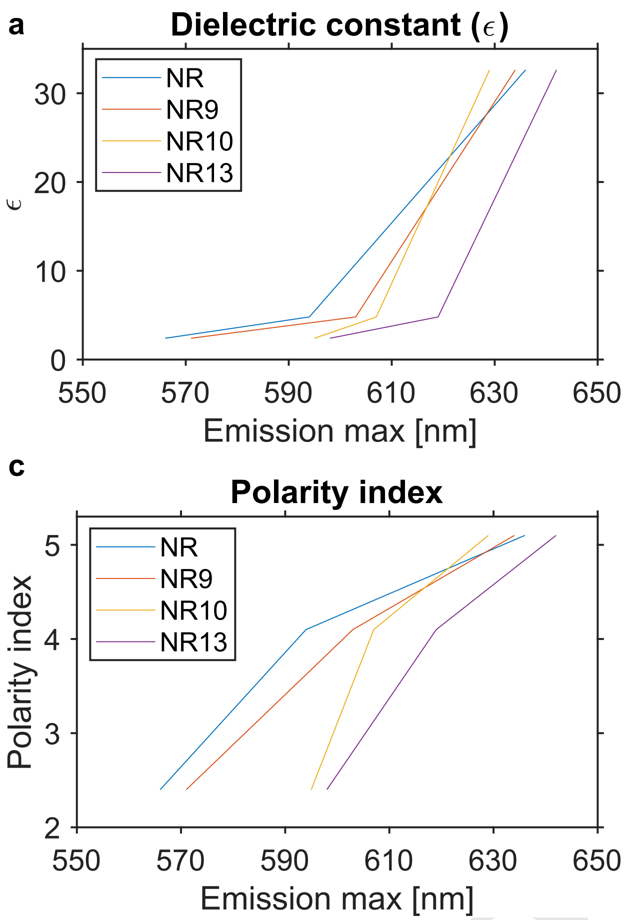


**Fig. S2.** Solvent properties as a function of emission max of Nile Red and the three analogs. Here, the dielectric constant(68) (a), the dipole moment(68) (b), the polarity index(68) (c), and the viscosity(69) (d) are plotted against the emission max. The emission max in solvents obtained from Hornum et al.(8) as shown in Tab. S1.

### Ratiometric study of the effect of cholesterol on the red emission of Nile Red, NR9, NR10, and NR13

Here, POPC liposomes with 0, 20, and 40 % cholesterol have been imaged in 2D STED and confocal and both signals have been fitted with a 2D Gaussian to extract the integrated intensity for each liposome. As expected, all four probes exhibit a green shift (lower ratio) upon increasing the cholesterol percentage in the liposomes because of the cholesterol-induced increase in membrane packing. This effect seems to be larger for the confocal data compared to the 2D STED data. Since we are capturing the confocal and the STED image simultaneously the two images also have the same pixel size, this means the confocal data will be largely over-sampled compared to the STED data, and there will be more pixels to define the fit, hence, the fit will be subjected to less noise, and the integrated intensity (and thereby the ratio) will be reflected with a higher accuracy.

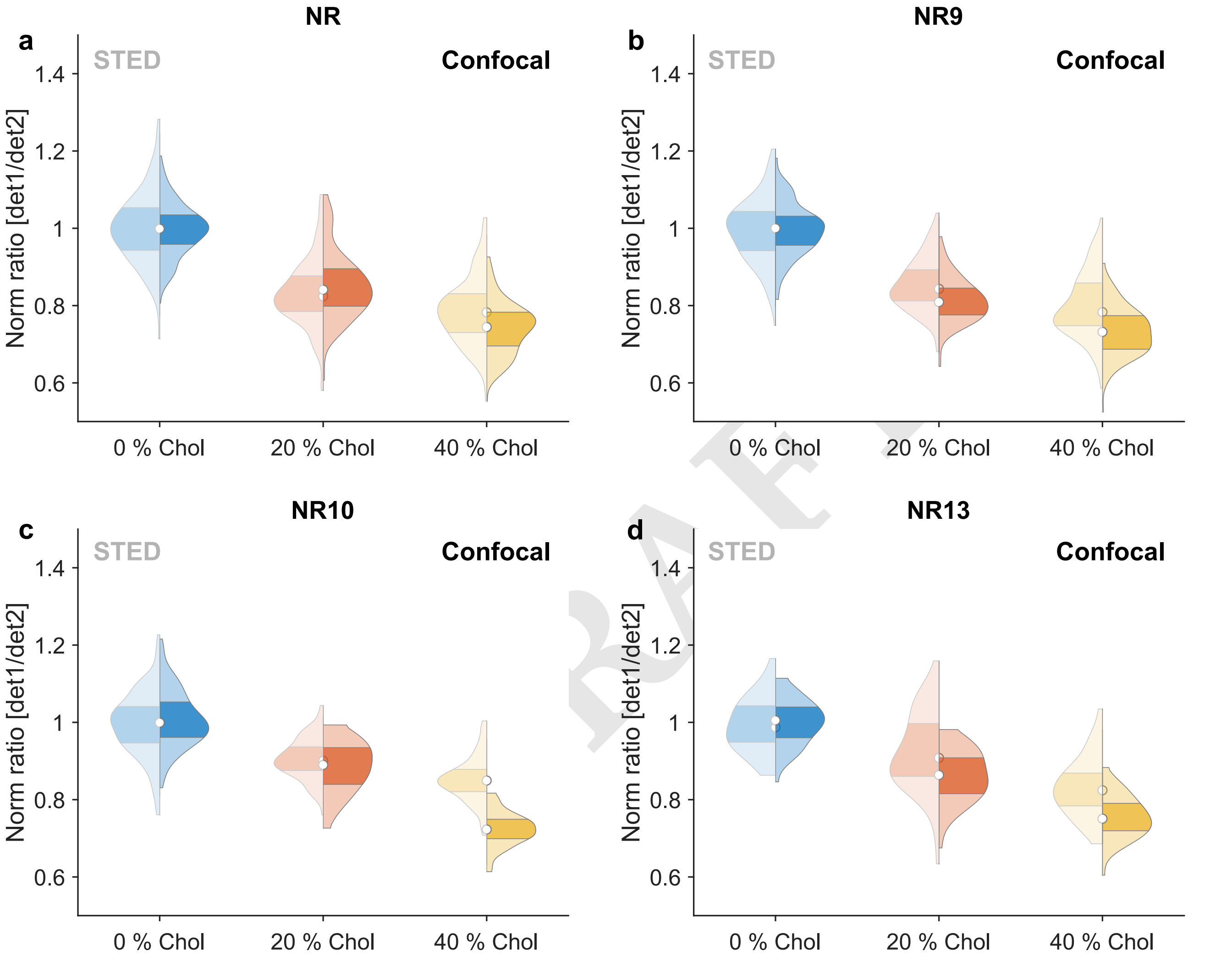


**Fig. S3.** Double-sided violin plots showing the STED (left) and the confocal (right) ratios measured for Nile Red (a), NR9 (b), NR10 (c), and NR13 (d) in POPC vesicles with 0, 20, and 40 % cholesterol, respectively. The ratios are normalized to the median ratio of POPC vesicles containing 0 % cholesterol, and due to noise outliers are removed from the data. Number of vesicles incorporated into the analysis for **Nile Red:** $N_{0\%Chol}$: 610, $N_{20\%Chol}$: 332, and $N_{40\%Chol}$: 500, **NR9:** $N_{0\%Chol}$: 234, $N_{20\%Chol}$: 233, and $N_{40\%Chol}$: 190, **NR10:** $N_{0\%Chol}$: 217, $N_{20\%Chol}$: 133, and $N_{40\%Chol}$: 120, and **NR13:** $N_{0\%Chol}$: 58, $N_{20\%Chol}$: 54, and $N_{40\%Chol}$: 49. Outliers are removed as values scaling more than three median absolute deviations.

To ease comparisons of the ratiometric studies between Nile Red and the three analogs, all ratios have been normalized to the median of the pure POPC vesicles. Then ratios >1 mean a less packed membrane and ratios <1 mean a more tightly packed membrane than POPC. Fig. S4 shows the raw ratios measured in vesicles and cells. Here, it is worth noticing, that the raw ratios measured in the vesicles and the cells are significantly higher than the ratios measured in MeOH, $CHCl_3$, and PhMe (Fig. S1f). This again shows that the spectral changes observed for the Nile Red probes in solvents and membranes should be compared only at a qualitative level.

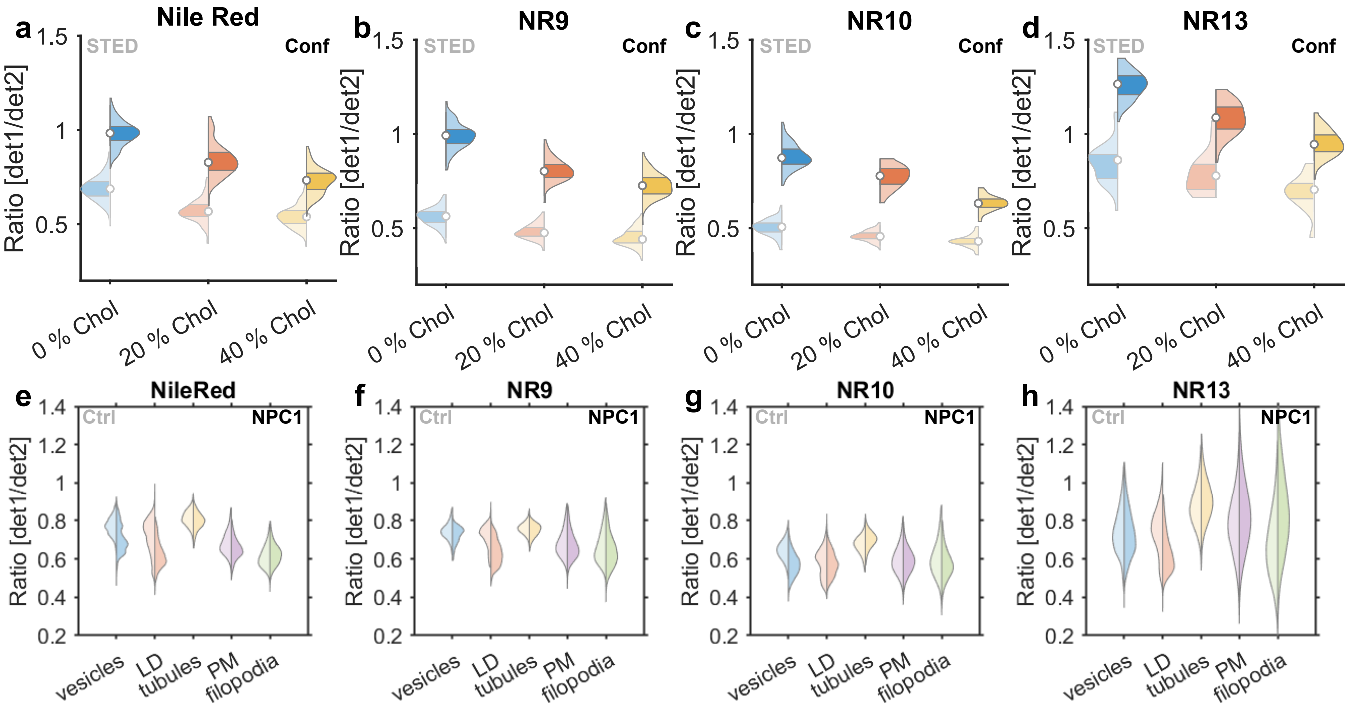


**Fig. S4.** Double-sided violin plots showing the non-normalized ratios (det1/det2) measured in vesicles and cells. For the vesicle measurements STED (left) and the confocal (right) non-normalized ratios measured for Nile Red (a), NR9 (b), NR10 (c), and NR13 (d) in POPC vesicles with 0, 20, and 40 % cholesterol, respectively. The same data as in Fig. S3, but not normalized to the median of pure POPC vesicles. (e-h) show double-sided violin plots which display the ratio for Nile Red (e), NR9 (f), NR10 (g), and NR13 (h) for different membrane compartments in control and NPC1 cells. This is the same data as in Fig. 5, but not normalized to the median of pure POPC vesicles. Outlier pixels are removed for all figures as values scaling more than three median absolute deviations

DRAFT

### Colocalization between DPH and green Nile Red fluorescence

To validate that the green signal from the Nile Red can function as a marker for the LDs the signal was colocalized with another LD marker, DPH. Fig. S5 show representative images of DPH staining (Fig. S5a), the green signal of Nile Red (Fig. S5b), and the red contribution of Nile Red (Fig. S5c). The Pearson coefficient is calculated between all three channels (Fig. S5d) showing a greater colocalization between DPH and the green signal of Nile Red, and a similar colocalization between the red signal of Nile Red and DPH or the green signal of Nile Red.

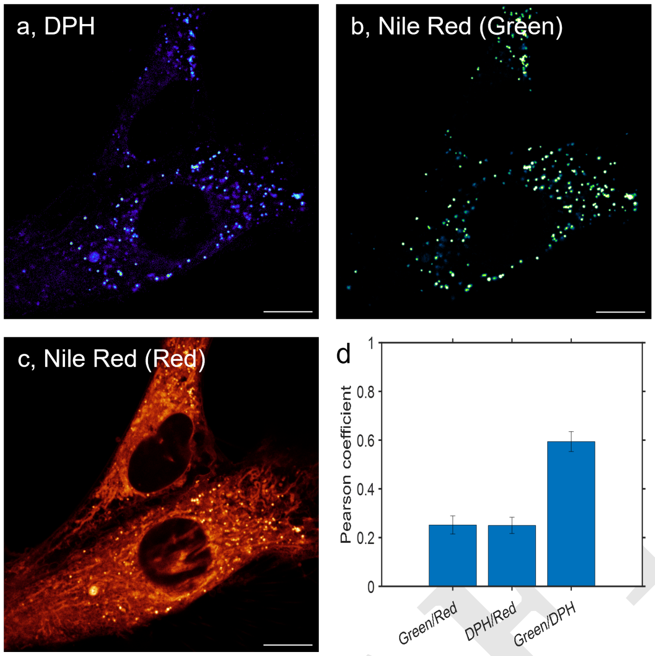


**Fig. S5.** Colocalization of lipid droplets with green emission of Nile Red and lipid droplet marker DPH. Confocal images of live control fibroblasts stained with DPH (a) and Nile Red (green emission, (b), and red emission, (c)). All scale bars are 10 µm. (d) Shows a bar chart of the mean Pearson coefficient calculated between the green and the red channel of Nile Red (Green/Red), DPH and the red channel (DPH/Red), and the green channel and DPH (Green/DPH) with the error bars showing the SEM ($N_{cells} = 9$).

DRAFT

### Masking sub-cellular membranes

To extract individual subcellular membrane compartments, these are drawn by hand on top of an image generated by summing the STED image measured in detector 1 and detector 2, as shown in Fig. S6a. A vesicle mask is drawn on top of circular structures and then divided into vesicles and LDs by the intensity signal found in the green emission. Here the lipid droplets of Fig. S6a are shown in Fig. S6b, and the remaining vesicles are shown in Fig. S6c, whereas the extracted tubular structures are shown in Fig. S6d.

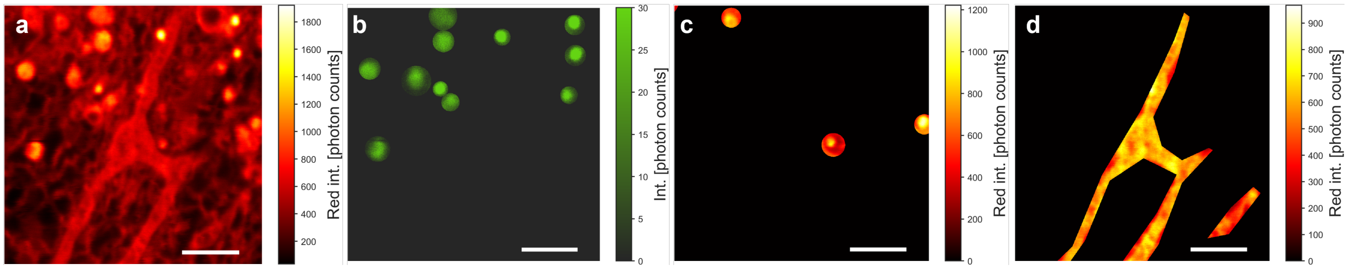


**Fig. S6.** Masking of different sub-cellular compartments in the area from Fig. 3a-d. (a) Summed intensity of Fig. 3a and b. (b) LD mask from Fig. 3c. (c) Vesicle mask based on a. (d) Tubules mask based on panel a. All scale bars 2 µm.

DRAFT

### Supplementary figures for the FLIM data

Lifetimes extracted by linearizing the FLIM data are plotted in the double-sided violin plot shown in Fig. S7 Nile Red, NR9, NR10, and NR13. We chose to linearize the function because the exponential fitting was too computationally demanding. Here, control fibroblasts are shown to the left and NPC1 cells are shown to the right. From this data, it seems that Nile Red derivatives in the NPC1 deficient cells have shorter lifetimes but also that their intensity was higher in NPC1 deficient cells compared to control cells. However, the data for the NPC1 deficient cells seemed to have a higher signal to noise. Therefore, we wanted to check if noisy data could generate longer lifetimes, and we generated a simple simulation of our data with a lifetime of 4 ns incorporating different noise levels.

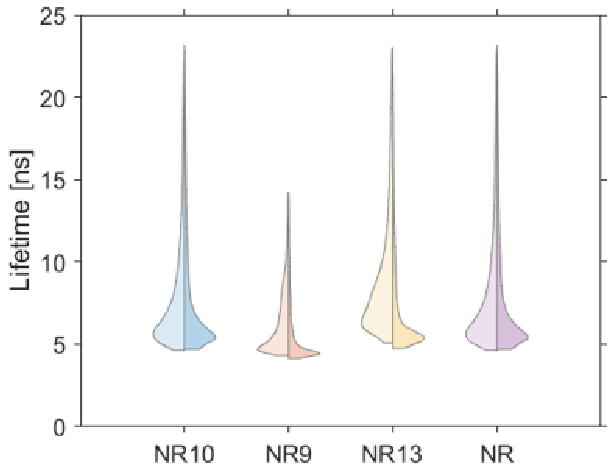


**Fig. S7.** Double-sided violin plot showing distributions of lifetimes from ctrl cells to the left and from NPC1 cells to the right obtained by transforming FLIM stacks with the natural logarithm and fitting each pixel with a linear regression. Lifetimes from control fibroblasts are shown to the left and NPC1 fibroblasts are shown to the right. Outlier pixels are removed as values scaling more than three median absolute deviations.

The simulated images are shown in Fig. S8a, where Fig. S8a.i shows the integrated FLIM intensity with no noise added, and then gradually more and more noise is added to each simulated image until Fig. S8a.v with the highest amount of noise. The data is then fitted with an exponential decay function and the lifetime is extracted. The lifetimes are shown in Fig. S8b. Then the data was log-transformed and the lifetimes were determined (Fig. S8c). When no noise is added the exponential and the linear fit performs equally well (Fig. S8b.i and c.i), and all the lifetimes extracted are 4 ns, which is the true simulated value. However, when we start to increase the noise it is observed that the extracted value of $\tau$ is also increasing. This is observed for the exponential fit (Fig. S8d), however, this tendency is much larger for the linearized function due to increased noise after log-transformation (Fig. S8e).

DRAFT

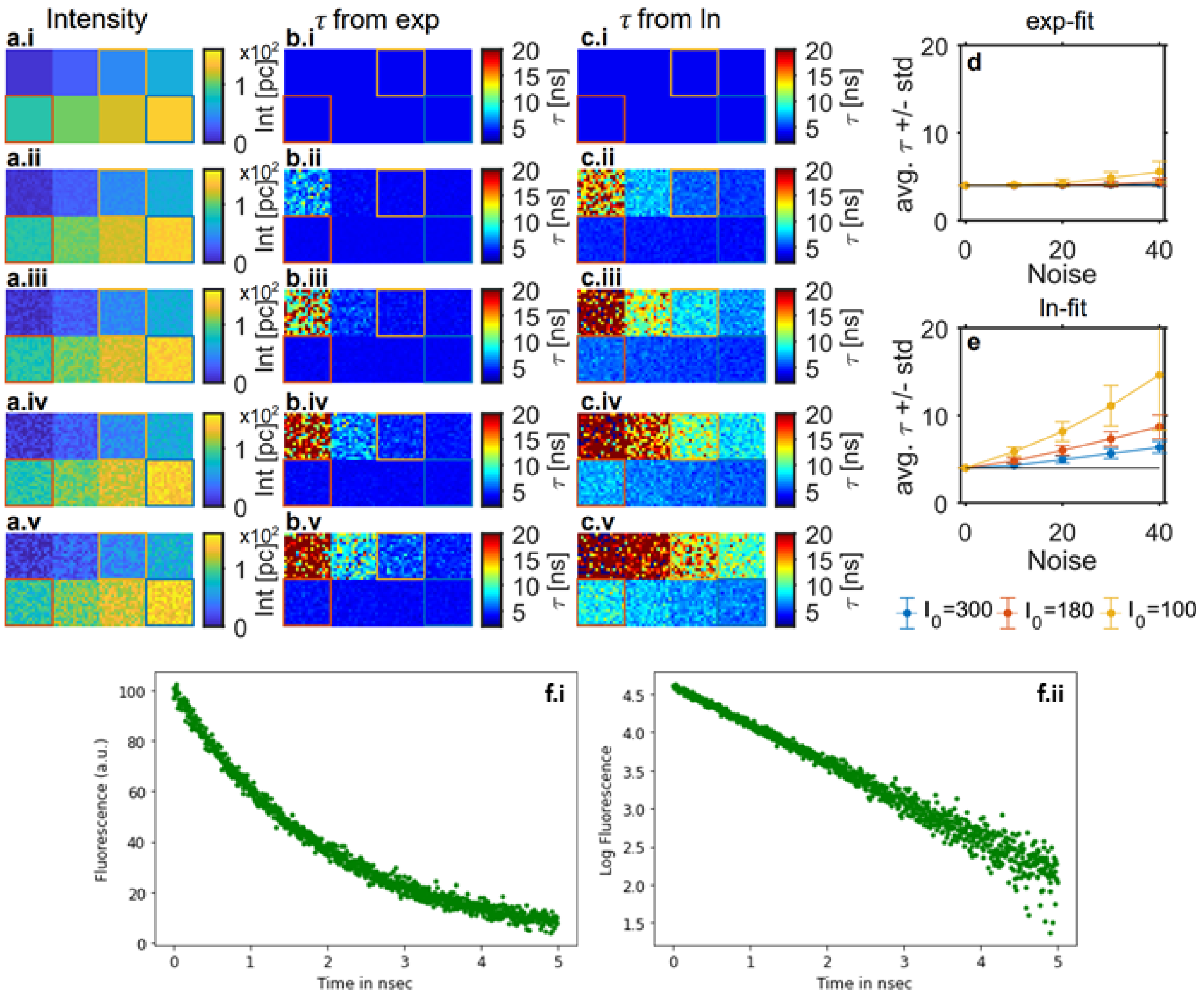


**Fig. S8.** Simulated FLIM stacks with the addition of different amounts of noise. (a.i)-(a.v) show the integrated intensity of simulated single exponential FLIM stacks from ±0-40 intensity units. (b.i)-(b.v) show the extracted lifetime values for each pixel extracted by a single exponential fit. (c.i)-(c.v) show the extracted lifetime values for each pixel extracted by linearization of the FLIM stack and fitting with linear regression. (d) shows the mean lifetime ($\tau$) obtained from the single exponential fit to the 16x16 pixel square of $I_0$=100 (yellow), 180 (red), and 300 (blue). (e) shows the mean lifetime ($\tau$) obtained from the linearized FLIM linear regression fit to the 16x16 pixel square of $I_0$=100 (yellow), 180 (red), and 300 (blue). (f) illustrates the effect of log transformation on noisy exponentially decaying intensity for a simulated fluorescence decay with additive Gaussian noise. One can see that noise gets redistributed in a non-linear manner, i.e., enhanced for longer times which results in ambiguity in slope estimation during linear regression.

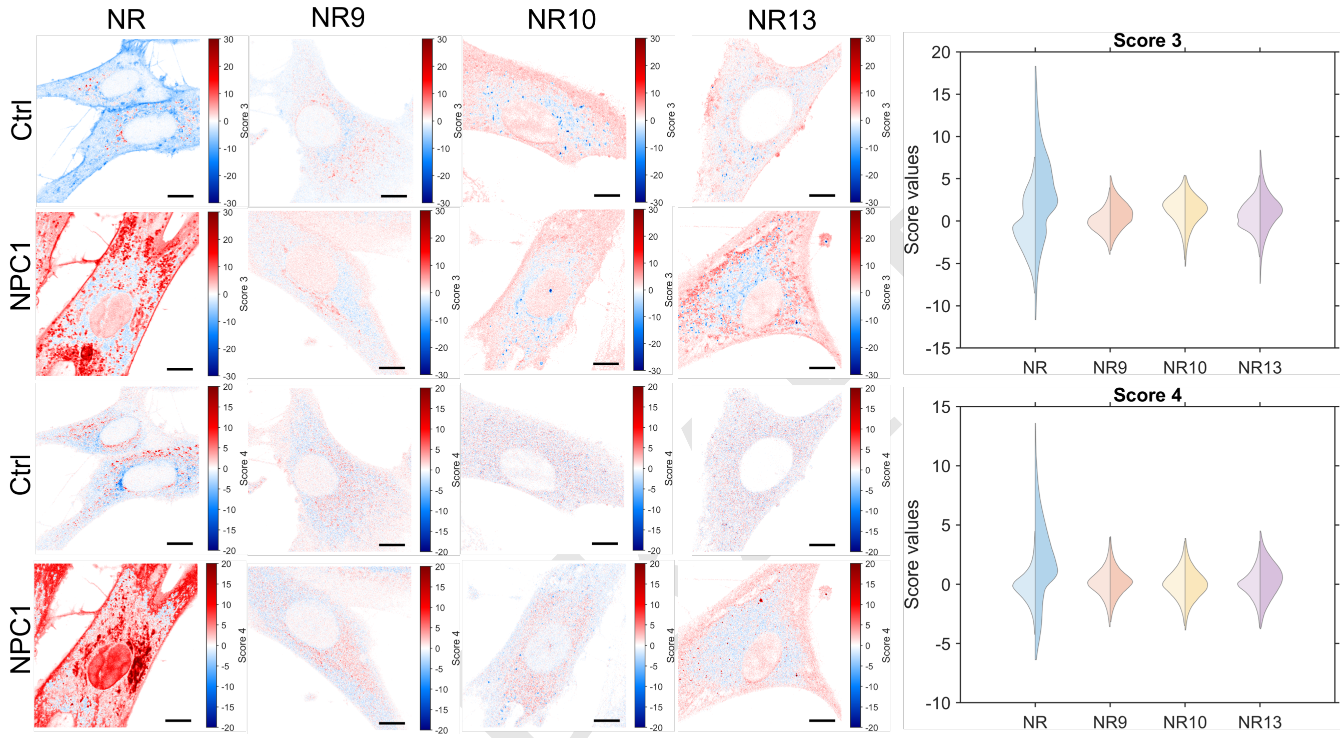


**Fig. S9.** Images for FLIM data generated by NC-PCA analysis for Nile Red, NR9, NR10, and NR13 labeled control and NPC1 cells. Here the estimates from score 3 and 4 are shown as images to the left and collected in double-sided violin plots to the right. The double-sided violin plots compare the distributions of score 3 (top) and score 4 (bottom) for control (left) and NPC1 (right) cells. Scale bars on the images are 10 µm. Outliers in the double-sided violin plots for score 3 and 4 are removed as values scaling more than three median absolute deviations.

### Supplementary MINFLUX analysis

**Single molecule binding events to vesicles.** To carry out single molecule tracking in isolated vesicles, we chose a dye concentration below the concentrations commonly used for fluorescence correlation spectroscopy (FCS). FCS is a good reference for our measurements, as it is based on quantification of fluctuations of fluorescence recorded from a confocal observation volume(70). Changes in the fluorescence fluctuations measured over time are then fitted with a temporal auto-correlation function. Since intensity fluctuations in FCS and other single-molecule techniques are based on the Poisson-distributed photon emission statistics, the extent of intensity fluctuations is inversely proportional to the mean intensity (71). This means that the more molecules are found in the detection volume, the smaller the relative fluctuations in fluorescence will be, therefore the number of molecules in the observation volumes need proper minimization normally between 0.1 and 1000, which corresponds to concentrations between 0.1 nM to $\approx$10 $\mu$M(70).

In our setup we are measuring single molecule binding to vesicles, taking advantage of Nile Red and the analogs' nature of being very bright when incorporated in a membrane (hydrophobic environment) and very dim when being in the buffer (hydrophilic environment). In Fig. S10a to the left the green fluorescence of a single vesicle is visualized by the incorporated NBD-PC lipid, used as a liposome marker. In the presence of 0.4 nM Nile Red, a confocal movie is recorded in the red channel, from which single frames are shown to the right. Since the microscope is scanning from left to right, and the binding and unbinding kinetics for Nile Red is faster than the scan of a single frame, binding and unbinding events of Nile Red to the vesicle will be shown as lines of higher and lower intensity, respectively(60). For a concentration of 0.4 nM Nile Red, there was almost always one or more binding events per frame. Therefore we chose to lower the concentration to 10 pM (Fig. S10b), where the frames from the movie are contrast adjusted to the same values as Fig. S10a.ii. Frame 4 in Fig. S10b.ii shows the background recordings of a single vesicle; these values would normally lie between 2-4 photon counts (pc). Since every pixel within the diffraction-limited spot of the vesicles is a measure of what is happening in the vesicle, extracting the maximum intensity per frame might be a good comparison for what is happening from 0.4 nM to 10 pM (Fig. S10c). Here, it is seen that for 10 pM the max intensity of

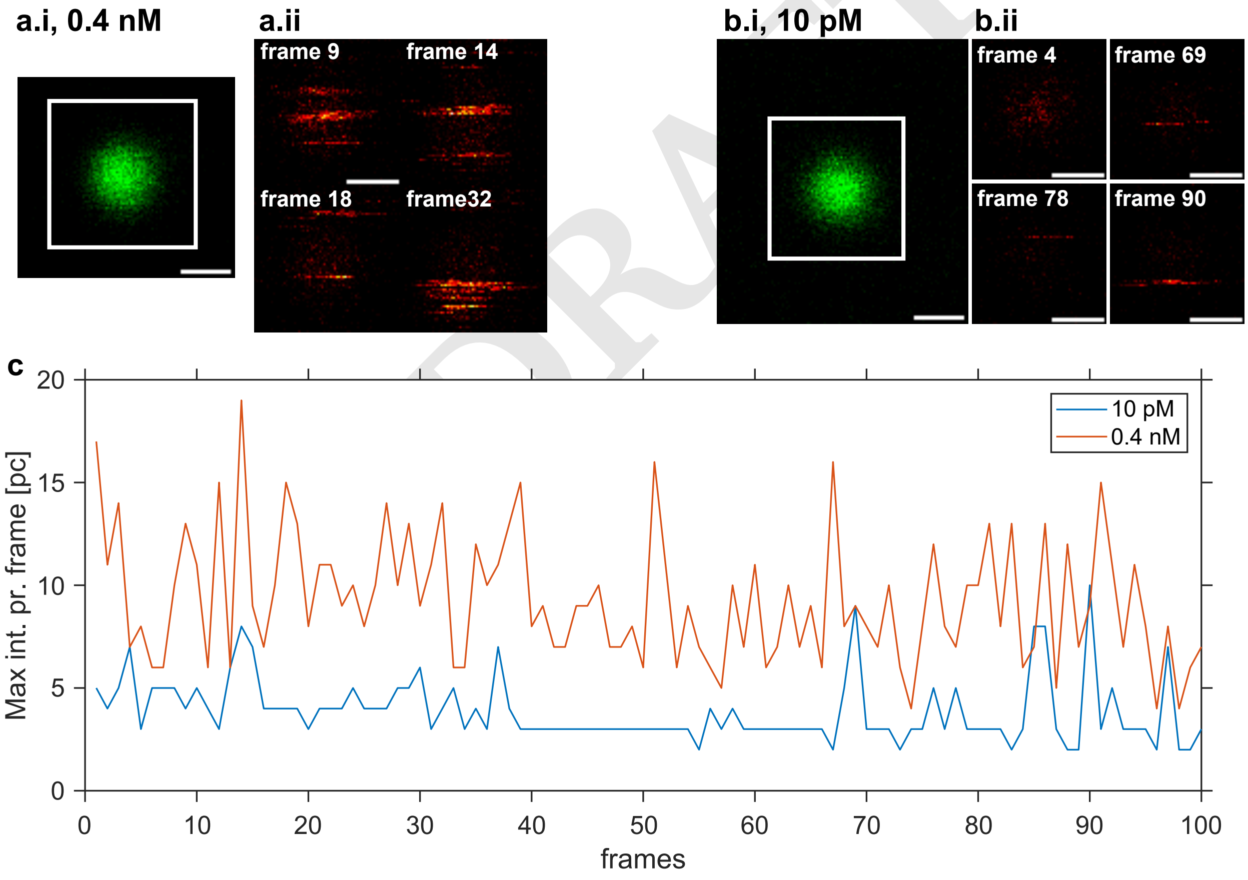


**Fig. S10.** Single molecule binding of Nile Red to single POPC vesicles labeled with 0.5 % NBD-PC. (a.i) Single POPC vesicle containing 0.5 mol% NBD-PC (ex. 488 nm, em. 500-550 nm). (a.ii) same vesicles as in a.i (white square) but imaged over time with a pixel dwell time of 10 $\mu$s in the presence of 0.4 nM Nile Red (ex. 561 nm, em. 580-630 nm). (b.i) Single POPC vesicle containing 0.5 mol% NBD-PC (ex. 488 nm, em. 500-550 nm). (b.ii) same vesicles as in b.i (white square) but imaged over time with a pixel dwell time of 10 $\mu$s in the presence of 10 pM Nile Red (ex. 561 nm, em. 580-630 nm). (c) shows the max intensity pr. frame plotted over the number of frames. The red plot shows the max intensity over time.

DRAFT

the frame lies close to the background intensity for single vesicles, between 2-4 pc, and for a molecule binding e.g. in frames 69 and 90 there is an intensity increase to around 10 pc. In a lot of frames acquired for 0.4 nM Nile Red, the maximal intensity is found around 10 pc or at 15 pc, which, minus the background, could correlate with twice the intensity, corresponding to two molecules binding the vesicle at this time.

**Diffusion coefficients of Nile Red, NR9, NR10, and NR13 of 2D MINFLUX tracks.** Single-molecule tracking carried out on a MINFLUX microscope varies from regular camera-based methods in two fundamental ways. One is the number of photons needed, since the tracking carried out here is more "active", as the microscope is continuously refocusing the beam onto the molecule, keeping the molecule as close to the center of the doughnut as possible. The other difference to camera-based imaging is that since Minflux tracking is based on the detection of photons, every localization step might take a different amount of time, which results in an uneven length of time intervals between molecule detection events (72). Therefore, to calculate the mean square displacement (msd), we calculated an average x and y position binned from time bins of 35 ms (similar size to the sliding window used by Balzarotti et al.(67)), and used these data to calculate the msd similarly to Lund et al.(73):

$$msd(t) = \frac{1}{N}\sum_{i=1}^{N}(x(i)-x(i+t))^2+(y(i)-y(i+t))^2 \tag{5}$$

Here, x and y are the spatial coordinates, $N$ is the number of frames, and $t$=$\delta t$ is the time increment. To analyze diffusion, the msd is fitted to the following model (73, 74):

$$msd(t) = 4\cdot D_{\alpha}\cdot t^{\alpha} \tag{6}$$

Where $D$ is the diffusion coefficient and $\alpha$ is the anomalous constant. If $0 < \alpha < 1$, molecules undergo subdiffusion, while $1 < \alpha < 2$ indicates superdiffusion. If $\alpha = 1$, molecules move by normal diffusion or Brownian motion (73, 74). Fig. S11a-d shows the msd calculated based on the single tracks obtained for Nile Red, NR9, NR10, and NR13. Here, the error bars show the standard deviation of the msd calculated for the individual molecules. All tracks have different lengths, depending on when a molecule binds, leaves, bleaches, or a second molecule enters a given vesicle. The time-averaged msd calculated according to Eq. 1 has an increasing error for larger time increments given that fewer data points enter the calculation. We also find that the time-averaged msd varies a lot between individual molecules. Since we have recorded a sufficient amount of molecule tracks lasting at least 1 s for all Nile Red derivatives, the diffusion coefficients and anomaly constants are determined from a fit of Eq. 6 to the first 1-s interval (Fig. S11e-h). Here, we find that NR13 and NR9 have $\alpha \approx 1$ which suggests a normal Brownian motion, whereas both Nile Red and NR10 have $\alpha$ values close to 2, which suggests a superdiffusive behavior. However, this behavior is not physically justified, and since we are recording longer tracks for both NR9 and NR13, which show Brownian motion (Fig. 7e), it seems that one needs to have long enough tracks for reliable estimation of molecular diffusion from MinFlux tracking data.

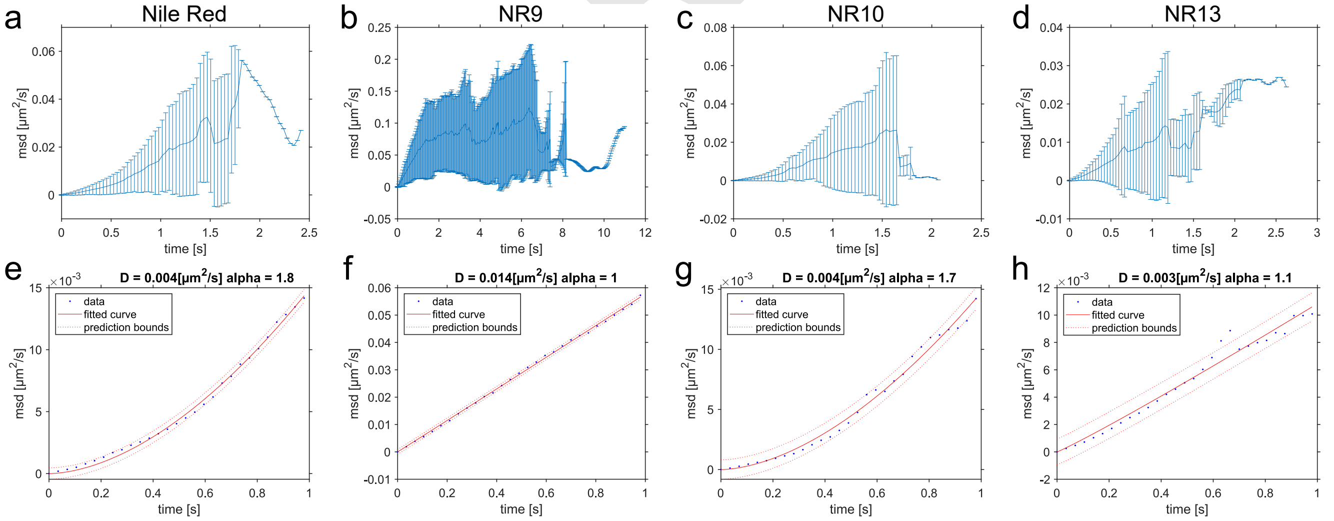


**Fig. S11.** Mean square displacement and diffusion coefficients calculated for Nile Red, NR9, NR10, and NR13. (a-d) msd calculated for Nile Red, NR9, NR10, and NR13 for a fixed time binning of 35 ms as in Eq. 5. (e-f) the first second of the time binned data shown in a-d fitted with Eq. 6 to calculate the $D$ and $\alpha$. Here, the blue dots show the data, the red line shows the fit, and the dotted red lines show the 95 % prediction bounds.

To investigate if the size of the time bins has a significant effect on $D$ and $\alpha$, we made the time bins smaller (10 and 20 ms) and one longer (100 ms), and fitted the msd with Eq. 6 (Fig. S12). Here, small and insignificant fluctuations were found for both $D$ and $\alpha$ when the time bins were increased in size, whereas the smaller time bins did not influence the results.

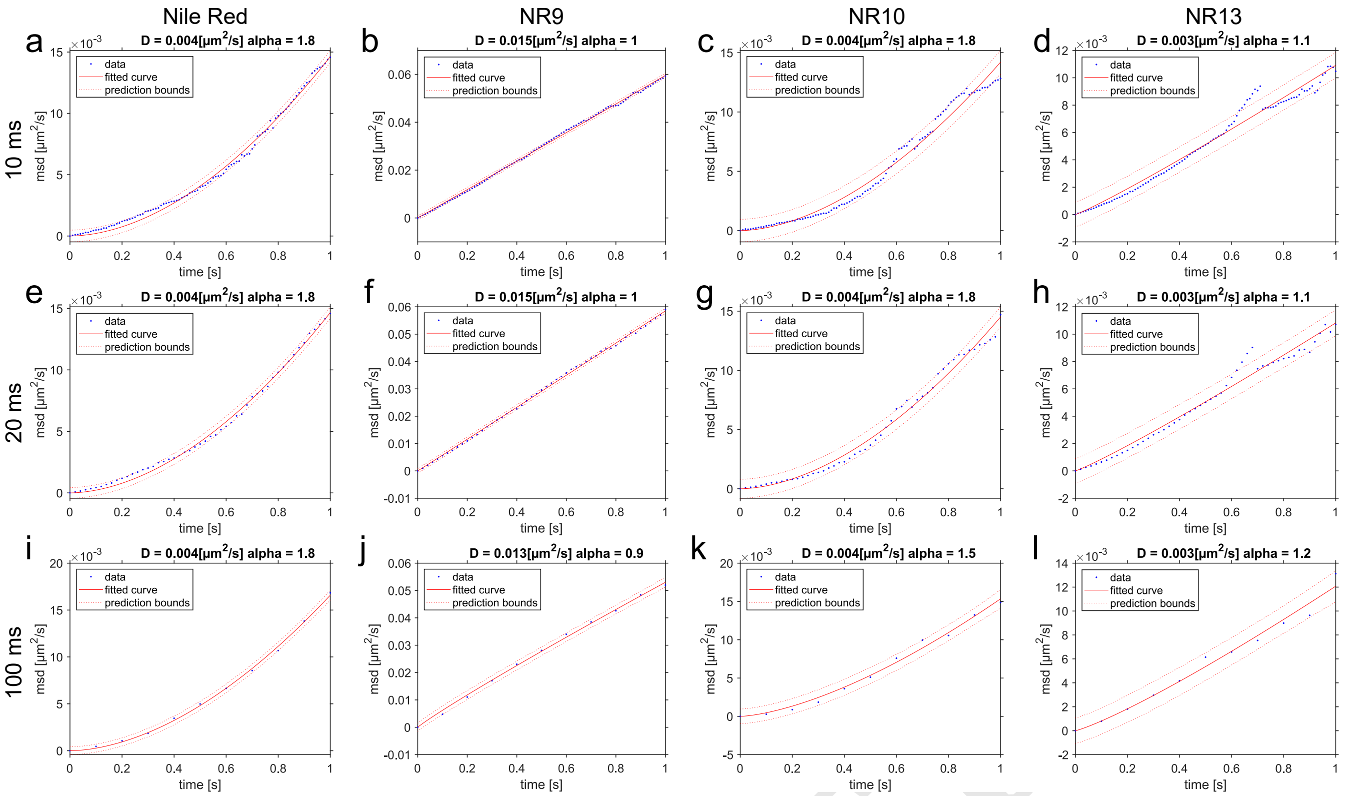


**Fig. S12.** Investigating the influence of different sizes of the time bins on the determination of the diffusion coefficient and the anomalous coefficient. Time bins of 10 ms (a-d), 20 ms (e-h), and 100 ms (i-l) for Nile Red, NR9, NR10, and NR13, with the calculated msd and a fit to Eq. 6 are shown and used to infer the values for $D$ and $\alpha$. Here, the blue dots show the data, the red line shows the fit, and the dotted red lines show the 95 % prediction bounds.

To investigate if the track length has an influence on $D$ and $\alpha$, we systematically cropped the length of the tracks entering the msd calculation. When the tracks are below 1 s long, the entire msd is fitted to Eq. 6, whereas when data points from tracks longer than 1 s are incorporated, it is only the first 1 s interval of the msd which is fitted. For the shorter tracks, there is a higher variability of the two parameters, whereas when the track lengths start to be longer than 2 s, there seems to be a stabilization of $\alpha$ and $D$, especially for NR9 and NR13, that also have longer tracks (Fig. S13). For Nile Red and NR10, most tracks are shorter than 1 s (Fig. 7e), which might also explain why $\alpha$ is not greatly affected by including the few longer tracks. This suggests, that if the track lengths of the molecules are too short (the majority of the tracks <1 s) in 2D MINFLUX tracking, apparent superdiffusion is observed in the msd analysis. The track length should be long enough for reliable inference of diffusion parameters.

DRAFT

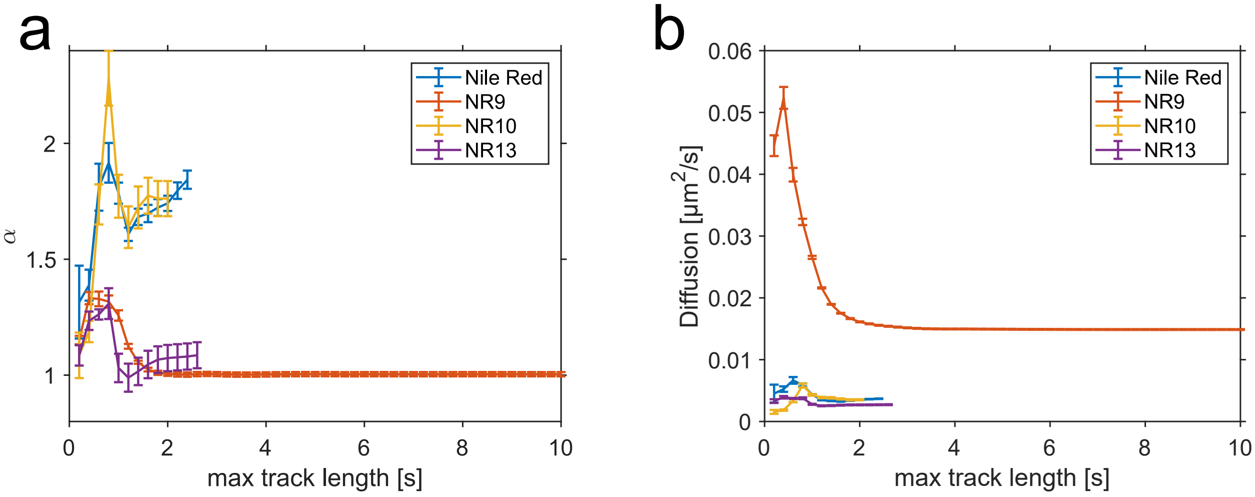


**Fig. S13.** *D* and $\alpha$ determined when varying the track lengths with time bins of 10 ms. (a) shows the extracted $\alpha$ over the maximum track length of the seconds. (b) shows *D* over the maximum track length. On both graphs, the error bars show the 95 % prediction bounds from the fit extracting $\alpha$ and *D*.

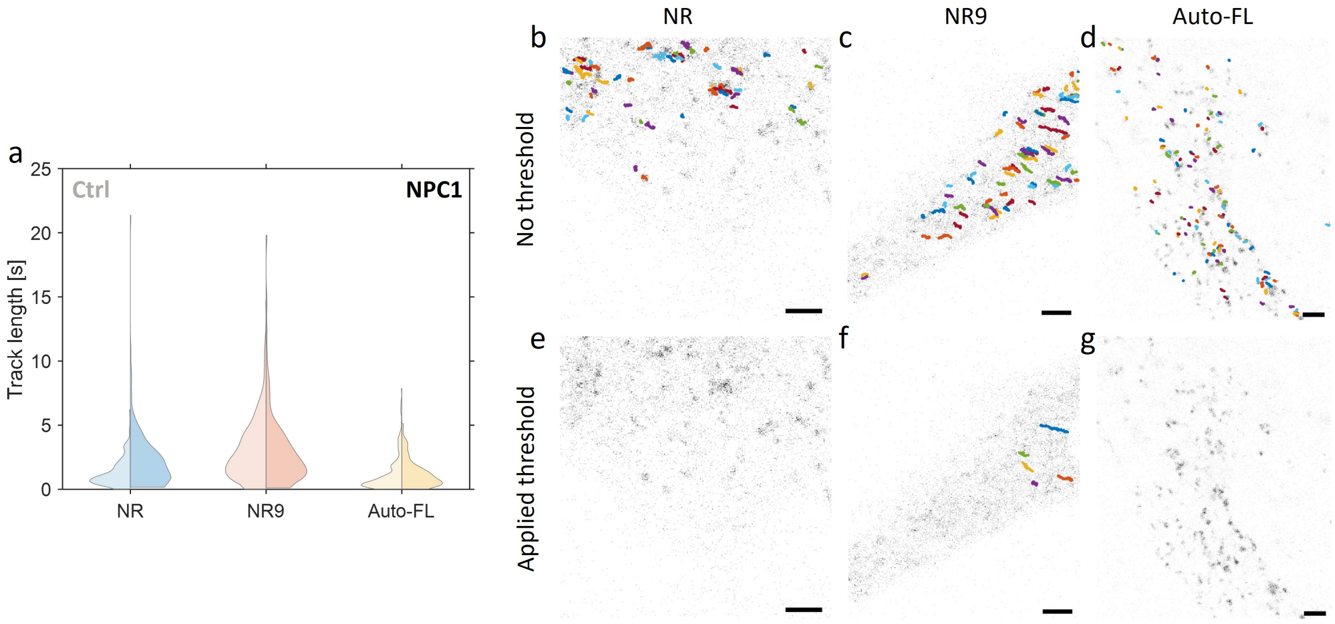


**Fig. S14. 2D MINFLUX tracking of Nile Red and NR9 in live ctrl and NPC1-deficient fibroblasts.** Trials with single-molecule tracking experiments of Nile Red and NR9 in control and NPC1 fibroblasts. (a) shows a double-sided violin plot of the distribution of single molecule track lengths of Nile Red, NR9, and autofluorescence (Auto-FL) in control fibroblasts (left) and NPC1 fibroblasts (right). The track length distribution for Nile Red in control fibroblasts contains tracks from 4 cells and 263 single molecule tracks, and the distribution for tracks of Nile Red in the NPC1 fibroblasts contains data from 5 cells and 206 single molecule tracks. The NR9 control fibroblast distribution contains tracks from 4 cells and 327 single molecule tracks, and the distribution for NR9 of the NPC1 fibroblasts contains data from 4 cells and 113 single molecule tracks. The Auto-FL control fibroblast distribution contains tracks from 4 cells and 191 single molecule tracks, and the Auto-FL distribution for the NPC1 fibroblasts contains data from 3 cells and 98 single molecule tracks. (b-d) show autofluorescence in the 560-nm channel of control fibroblasts in grayscale, and individual single-molecule MINFLUX tracks on top for 10 pM of Nile Red (b), 10 pM of NR9 (c), and Auto-FL (d). (e-f) show the same images as in b-d, but only with the single molecule tracks which were longer than the ones observed in the Auto-FL. All scale bars are 2.5 μm.
